# Concentration-Dependent Mutational Scanning Probes the Cellular Folding Landscape of *α*-Synuclein in Yeast

**DOI:** 10.1101/2025.08.29.672156

**Authors:** Daeun Noh, Robert W. Newberry

## Abstract

The misfolding and aggregation of α-synuclein is a central molecular event in the etiology of Parkinson’s disease and related disorders. α-Synuclein misfolding and pathology are both concentration-dependent, but it is not clear precisely how changes in concentration alter the folding landscape within cells. Whereas most conventional structural biology approaches offer limited resolution in living systems, deep mutational scanning can offer insight into the folding state of a protein in living cells, and we apply this method to probe concentration-dependent changes in the folding of α-synuclein in a popular yeast model of pathology. We discover that at a wide range of cellular concentrations, α-synuclein is highly biased toward formation of a membrane-bound amphiphilic helix that imparts toxicity. Population of this toxic state can be disrupted by mutations that reduce membrane affinity, which shift the folding equilibrium away from the membrane-bound state. Reduced-affinity variants exhibit distinct sensitivity to concentration relative to variants with WT-like affinity, likely because these variants are expressed at concentrations closer to their dissociation constant for membrane binding. These results show how mutational scanning can provide high-resolution insights into the folding landscape of proteins in living cells, which is likely to be of special utility for studying proteins that misfolding and/or aggregate.

**Impact Statement:** Protein misfolding is often concentration-dependent, but studying concentration-dependent changes in folding in living cells is challenging. By using high-throughput mutagenesis, we reveal changes in the population of toxic conformations of the Parkinson’s-associated protein α-synuclein. We discover that in a yeast model of pathology, α-synuclein is highly biased toward membrane binding, which in turn disrupts cellular homeostasis.

## Introduction

α-Synuclein is an abundant neuronal protein implicated in the physiological trafficking of synaptic vesicles during neurotransmission, as well as the pathological cellular degeneration in Parkinson’s and related diseases. Understanding the role of α-synuclein in these processes has been complicated by its wide conformational repertoire (Bendor et al., 2013; Lashuel et al., 2013; So & Watts, 2023). For example, purified α-synuclein is intrinsically disordered *in vitro* (Eliezer et al., 2001; Weinreb et al., 1996) and can retain that disorder in living cells (Theillet et al., 2016). Upon association with lipid membranes, α-synuclein can adopt a variety of helical conformations with amphiphilic character that enable membrane adsorption via seven imperfectly repeated 11-residue membrane-binding segments (Davidson et al., 1998; Bussell & Eliezer, 2003; Jao et al., 2004; Fusco et al., 2014; Dettmer et al., 2015; Fusco et al., 2016). These structures are likely essential for its physiological roles in neurotransmission (Davidson et al., 1998; Fusco et al., 2014), but dysregulation of these structures is also implicated in disease (Burré et al., 2014; Mansueto et al., 2023). For example, excessive population of the membrane-bound state can disrupt membrane integrity and vesicular trafficking (Cooper et al., 2006; Busch et al., 2014; Fusco et al., 2017; Kim et al., 2021).

α-Synuclein can also misfold into a wide variety of structures that feature varying degrees of polymerization and/or secondary structure, particularly β-sheet. These species range in size from oligomers of a few monomers to amyloid fibrils that can be hundreds of nanometers in length and incorporate thousands of monomers. These misfolded states are highly polymorphic and can differ in the fold of individual monomers, the packing/stacking of monomers against one another, and the ultrastructure of the resulting aggregate, all of which can affect the pathogenicity of the material (Knowles et al., 2014; Gallardo et.al., 2020; Louros et al., 2023; Milchberg et al., 2025). Misfolded states often take on new activities that disrupt cellular homeostasis, such as by distracting proteostasis machinery, disrupting vesicular trafficking, and compromising membrane integrity. (Lashuel, 2005; Chiti & Dobson, 2017). As a result, misfolded α-synuclein is implicated as a central player in the etiology of Parkinson’s and related diseases (Lashuel et al., 2013; Wong & Krainc, 2017; Fares et al., 2021; Calabresi et al., 2023).

Ongoing interest in understanding and treating Parkinson’s and related diseases is complicated by α-synuclein’s diverse conformational repertoire and associated mechanisms of pathology (Villar-Piqué et al., 2015; Li & Liu, 2022; So & Watts, 2023). One unifying feature of α-synuclein toxicity is its concentration dependence, which is observed both in humans, as well as many common laboratory models. For example, the duplication and triplication of the *SNCA* gene that encodes α-synuclein have been observed in families with inherited, early-onset Parkinson’s (Bradbury, 2003; Ibáñez et al., 2004; Konno et al., 2016; Singleton et al., 2003), indicating the dose-dependent nature of synuclein toxicity in humans. Increased gene dosage of α-synuclein in microbes, cultured mammalian cells, and transgenic mice also reliably impart features of Parkinson’s (Outeiro & Lindquist, 2003; Luk et al., 2012; Chung et al., 2013).

Changes in α-synuclein concentration can affect the folding landscape and subsequent toxicity in a variety of ways. First, α-synuclein aggregation is a concentration-dependent process in both *in vitro* and *in vivo* ( Iljina et al., 2016 ; Perrino et al., 2019; Wood et al., 1999). Increased concentration increases the likelihood of initial nucleation and accelerates various mechanisms by which aggregation propagates to subsume additional monomers (Buell et al., 2014; Meisl et al., 2016; Afitska et al., 2019). In turn, increased aggregation can amplify associated mechanisms of pathology.

Increased α-synuclein concentration can also increase the frequency of aberrant interactions with other cellular components. For example, increases in α-synuclein concentration increase the membrane-bound population *in vitro* and in cells (Outeiro & Lindquist, 2003; Galvagnion et al., 2015). Excessive concentrations of this population are associated with membrane permeabilization, global protein-trafficking stress, and impaired import of nuclear-encoded mitochondrial proteins (Cooper et al., 2006; Tosatto et al., 2012; Di Maio et al., 2016). Moreover, excessive membrane binding can in turn drive subsequent self-association and aggregation (Galvagnion et al., 2015; Makasewicz, et al, 2024). Adsorption of α-synuclein from a three-dimensional volume onto a two-dimensional surface effectively increases the local concentration, thereby driving aggregation. Electron microscopy of Lewy bodies, the pathological hallmark of Parkinson’s, shows high densities of aggregated α-synuclein and membrane-bound structures, implicating both membrane-dependent and -independent mechanisms of α-synuclein aggregation and toxicity (Shahmoradian et al., 2019).

Furthermore, recent studies have discovered concentration-dependent conformational changes in membrane-bound α-synuclein. Specifically, at lower concentrations, EPR spectroscopy shows an extended helical conformation with the N-terminal region and part of the NAC region bound to the membrane (Georgieva et al., 2008; Jao et al., 2008), whereas a partially released, upright conformation was observed at higher concentration (Roeters et al., 2023). Release of the aggregation-prone NAC region could further accelerate membrane-dependent (Snead & Eliezer, 2019) and exacerbate the effect of concentration on α-synuclein toxicity.

Concentration-dependent changes in α-synuclein folding, binding, and aggregation are therefore of great importance for understanding the mechanisms of toxicity. Critically, we must understand which of these various concentration-dependent processes are operative in different cellular environments. Doing so would allow the community to employ appropriate models for studying specific molecular processes and possible interventions that disrupt then. Whereas cellular studies to date have identified phenotypic changes in response to changing α-synuclein concentration (Delenclos et al., 2019), much less is known about the conformations of α-synuclein that mediate those concentration-dependent phenotypes.

Unfortunately, probing protein conformation in living cells with high resolution remains challenging, thereby thwarting efforts to understand which conformations of α-synuclein contribute to specific pathogenic phenotypes, including those that are concentration-dependent. In-cell NMR has offered some inroads into the conformation of α-synuclein in mammalian cells (Binolfi et al., 2016; Theillet et al., 2016; Kragelj et al., 2025; Dumarieh et al., 2025), but limitations in instrument sensitivity, isotope-labeling schemes, and pulse sequences restrict the species that can be detected and make concentration-dependent measurements challenging.

A promising alternative is to infer structural features of proteins from the effect of mutations on the activity of the protein. Doing so requires measuring the effects of mutations throughout the protein, which is made possible by advances in high-throughput mutagenesis, such as deep mutational scanning (DMS) (Fowler & Fields, 2014; Wei & Li, 2023). In DMS, a library of protein variants is screened for activity in a high-through selection to create a comprehensive mapping of protein genotype to cellular phenotype. The resulting sequence–activity landscapes highlight a variety of biochemical features of the active state of the protein, including elements of secondary and tertiary structure (Rollins et al., 2019; Schmiedel & Lehner, 2019; Tsuboyama et al., 2023). Indeed, DMS has been employed to unravel molecular determinants of protein aggregation and structures of various misfolded proteins including Aβ, α-synuclein, and TAR DNA-binding protein 43 (TDP-43) (Bolognesi et al., 2019; Gray et al., 2019; Newberry et al., 2020; Seuma et al., 2022; Chlebowicz et al., 2023; Thompson et al., 2025; Arutyunyan, et al., 2025). Furthermore, a recent study showed that DMS data have been used to train a neural network model for predicting the aggregation of proteins from their sequences (Thompson et al., 2025).

For example, we previously applied DMS to probe the pathology of α-synuclein in yeast, a minimal cellular model (Newberry et al., 2020). In that system, mutations that disrupt the toxicity of the protein consistently destabilize interactions between α-synuclein’s amphiphilic helix and lipid membranes (Bodner et al., 2009; Fusco et al., 2014). For example, proline residues disrupt toxicity by interfering with helix formation, while polar substitutions on the hydrophobic face of the helix disrupt toxicity by reducing membrane affinity. In fact, a simple thermodynamic model of this structure was sufficient to predict the phenotypes observed for α-synuclein variants in yeast cells (Newberry et al., 2020). This structure is consistent with cellular effects of α-synuclein expression in yeast, which primarily perturbs membrane-dependent processes (Outeiro & Lindquist, 2003; Cooper et al., 2006; Gitler et al., 2008; Yeger-Lotem et al., 2009). DMS is therefore a powerful approach to understanding the pathogenic states of proteins in high resolution.

In this study, we apply DMS to probe how changes in α-synuclein concentration contribute to cellular pathology by altering the population(s) of its possible structural states. We are particularly interested in two fundamental questions. First, do changes in concentration affect the structure of the pathogenic species? This could occur through several possible mechanisms. On one hand, some species might only accumulate and/or become pathogenic at threshold concentrations that exceed the proteostasis capacity of the cell. For example, the amphiphilic helix might dominate toxicity at high concentrations, whereas at lower concentrations, toxicity could be dominated by other species that might be less prevalent but more successful at evading proteostasis. Alternatively, self-interactions at high concentrations might drive conformational changes that range from relatively subtle (e.g., vertical vs horizontal poses of the membrane-bound helix) to dramatic (e.g., amyloid nucleation of disordered/helical monomers). Changes in structure and dynamics should be evident in the set of mutations that disrupt proteotoxicity. For example, transition from α-helical to β-sheet structure should change the periodicity in mutations that disrupt toxicity. We are especially interested in this question given the high levels of expression we employed in previous DMS of α-synuclein toxicity (Newberry et al., 2020; Newberry et al., 2020a).

Second, how do changes in concentration shift the populations between coexisting conformational states in living cells? Previous DMS studies of α-synuclein at high expression levels only identified mutations that disrupt toxicity, rather than enhancing it. We hypothesize that the lack of identified enhancers results from high expression heavily biasing the conformation equilibrium toward the toxic conformational state. We therefore predict that changes in concentration might shift the conformational equilibrium. For example, a decrease in concentration could shift the equilibrium away from the membrane-bound state toward the disordered, unbound state. Alternatively, decreasing concentration could simply reduce the population of the pathogenic, membrane-bound state without shifting equilibrium away from membrane binding. The concentration-dependence of mutational effects in DMS should distinguish between these possibilities. Specifically, a simple reduction in the membrane-bound population should manifest as reduced sensitivity to mutation overall, whereas a shift toward the unbound population might result in the emergence of mutations that enhance toxicity by driving equilibrium back toward the toxic state, which would be especially significant for understanding determinants of α-synuclein toxicity given that our previous DMS studies failed to identify any such enhancers.

## Results

### Experimental design

Despite a lack of synuclein homologs, budding yeast (*Saccharomyces cerevisiae*) has become a powerful cellular model for studying α-synuclein toxicity and aggregation (Khurana & Lindquist, 2010). Heterologous expression of α-synuclein in yeast recapitulates several key features of its pathology in human neurons, including membrane-binding, aggregation, and dose-dependent toxicity. (Outeiro & Lindquist, 2003; Sampaio-Marques et al., 2019). α-Synuclein puncta in yeast incorporate ER and Golgi-associated vesicles (Soper et al., 2008), mirroring the Lewy bodies seen in PD patients (Dettmer et al., 2017; Gitler et al., 2008; Shahmoradian et al., 2019; Trinkaus et al., 2021). Furthermore, the yeast model has been successful used to discover both chemical and genetic modifiers of α-synuclein toxicity, which have been subsequently validated in neuronal and animal studies (Chung et al., 2013; Cooper et al., 2006; Fanning et al., 2019; Gitler et al., 2009; Khurana et al., 2017; Sangkaew et al., 2022; Yeger-Lotem et al., 2009). Previous studies of α-synuclein in yeast have shown predominance of the N-terminally acetylated species due to robust expression of the N-terminal acetylation machinery, NatB (Vamvaca, et al., 2009; Zabrocki et al., 2008). We therefore expect that our results reflect the behavior of the acetylated protein.

Yeast also offers convenient strategies for controlling protein expression through titration of inducible promoters. In previous DMS studies (Newberry et al., 2020), we induced α-synuclein expression from the GAL promoter at a galactose concentration of 1% (w/v). Given this high level of expression, we were concerned that we had driven the α-synuclein population toward states that are not relevant at lower expression levels. Indeed, our previous studies were unable to identify mutations that enhance α-synuclein toxicity at this concentration, suggesting that we had driven expression toward a toxicity ceiling.

We therefore sought to identify conditions that would induce lower levels of α-synuclein expression. To do so, we compared the growth of yeast strains expressing α-synuclein-GFP to strains expressing just GFP at a variety of inducer concentrations (Figure S1). As expected, decreasing the concentration of the inducer reduced the growth defect of yeast expressing α-synuclein relative to those expressing GFP down to a concentration of 0.001% (w/v), where no α-synuclein toxicity was detected. We confirmed α-synuclein expression at inducer concentrations above 0.001% (w/v) by tracking fluorescence of the GFP tag. Microscopy of induced cells showed greater GFP fluorescence than uninduced samples (Figure S2). Based on these results, we selected four different inducer concentrations for subsequent DMS experiments (i.e., 0.001%, 0.01%, 0.1%, and 1% [w/v]).

A yeast library expressing 2,600 of the 2,660 possible single missense variants of α-synuclein was developed previously (Newberry et al., 2020; Newberry et al., 2020a). In this library, each variant was associated with approx. 12 random DNA barcodes to expedite downstream analysis. This library was induced at concentrations described above, and samples were collected from the library over time. Deep sequencing of the DNA barcodes recovered from these samples reveal changes in the frequency of each variant over time, which we interpret as a fitness score (i.e., a measure of the toxicity of that variant) (Table S1). Each selection experiment was performed in duplicate, and duplicates at all four inducer concentrations showed strong agreement (Figure S3). By comparing these fitness scores for each variant across concentrations, we were able to probe changes in folding landscape of α-synuclein in these living cells (Figure 1A).

**Figure 1.**
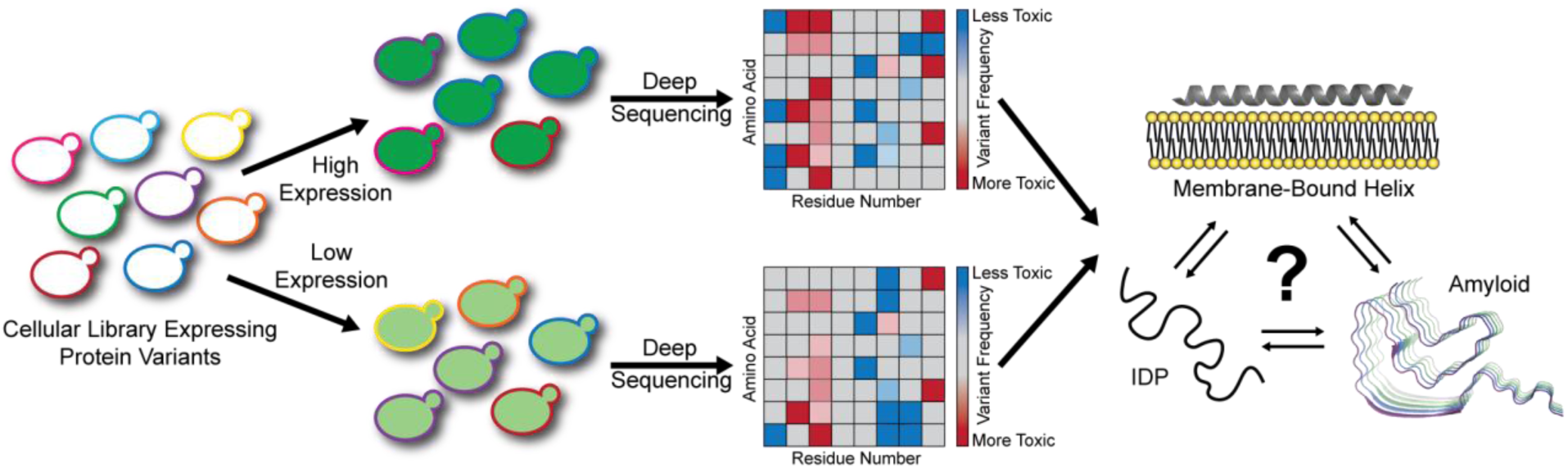
Mutational approach to probe *α*-synuclein conformational states at different protein expression levels. An α-synuclein library including 2600 single missense variants was expressed in yeast. The protein was induced to different expression levels by varying the concentration of the chemical inducer (i.e., galactose), resulting in distinct selective pressures that differentially alter the relative frequencies of each variant in the population over time. The frequencies of each variant were counted using deep sequencing and converted to a fitness score that reflects their toxicity relative to WT. The resulting sequence–toxicity landscape reveals key structural features of the protein in its toxic state, thereby reporting on changes to the conformational landscape as a function of protein concentration.

### Conservation of structure in the toxic conformation of α-synuclein across expression levels

To identify possible changes in α-synuclein’s toxic conformation as a function of concentration, we first mapped the fitness scores of all 2600 single amino-acid substitutions at each inducer concentration (Figure 2A-D, Table S2). Lower inducer concentrations result in weaker selective pressures and by extension, a condensed range of fitness scores. To capture that change in the dynamic range, we have plotted all fitness scores in Figure 2 on the same scale so that the effect of mutations can be compared between datasets. Within each dataset, the average fitness score of barcodes mapping to WT is set equal to zero and all other fitness scales are adjusted accordingly. To validate our findings from the high-throughput assay, we selected variants that sample the full range of fitness scores for individual retest at each inducer concentration; this data is now included in the supplementary information as Figure S4. As expected, each mutation causes a smaller change in growth rate at lower inducer concentrations, but the rank order of toxicity remains the same across inducer concentrations.

**Figure 2.**
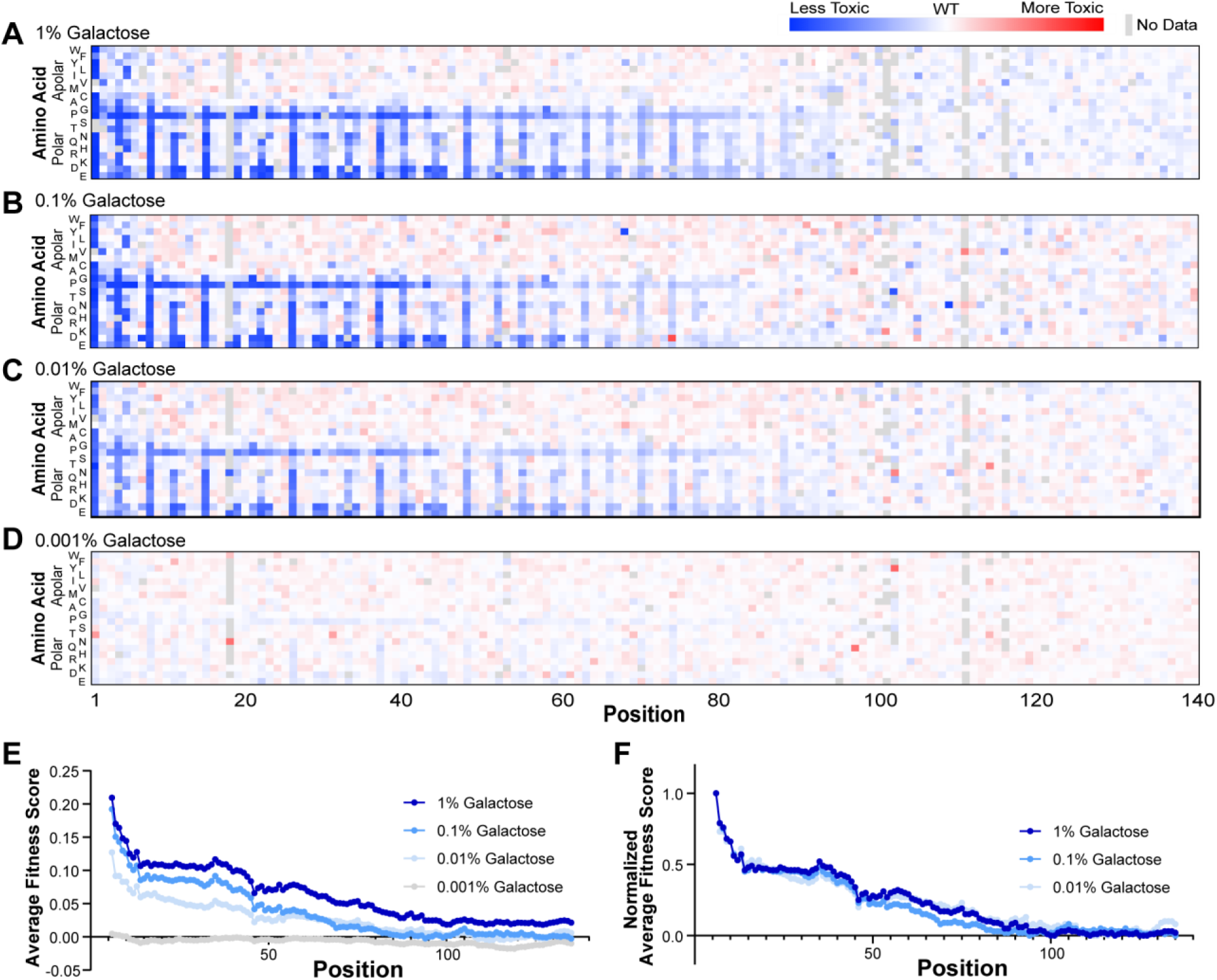
Fitness scores of *α*-synuclein variants at different protein expression levels. Fitness scores were calculated following selection of α-synuclein variants based on toxicity induced by galactose concentrations of **(A)** 1%, **(B)** 0.1%, **(C)** 0.01%, and **(D)** 0.001% (w/v). **(E)** Average fitness scores over an 11-residue window centered at each position in α-synuclein and **(F)** the same fitness scores as (E) normalized to the maximum fitness score at each inducer concentration.

Remarkably, we observed robust conservation in the pattern of mutations that disrupts α-synuclein toxicity in yeast. Specifically, at all expression levels, proline substitutions in the first ∼90 residues consistently reduced toxicity. The same was true for incorporation of polar amino acids onto the hydrophobic face of the amphiphilic helix, manifesting as a strict 3.67-residue periodicity in the effects of polar substitutions on toxicity (Figure S5). Moreover, the effects of mutations associated with human synucleinopathies (Table S6, Figure S6) were also well-correlated with their effects on the stability of the membrane-bound helix. Effects of these mutations on other processes (e.g., amyloidogenesis) correlate less well with effects on yeast toxicity and the concentration dependence thereof. For example, the A53T mutation, which generally increases the aggregation rate of purified α-synuclein *in vitro* (Conway et al., 1998; Narhi et al., 1999), causes little change in yeast toxicity across the concentrations sampled herein. This provides further support for the membrane-bound helix as the driver of toxicity in this yeast model.

As concentration decreases, the effect of each mutation generally decreases in magnitude, but the pattern of disrupting mutations remains consistent. Because the overall pattern of mutations that disrupts α-synuclein toxicity in yeast is unchanged as a function of concentration (Figure S7), we conclude that there are no gross changes in structure, either due to self-association or masking of conformations resolved by proteostasis. The robustness of this toxic conformational state demonstrates that it is not an artifact of high expression level and instead further emphasizes its centrality in this cellular model.

To evaluate more subtle changes in the structure and/or dynamics of the amphiphilic helix caused by changes in expression level, we calculated the average effect of substitutions in sliding 11-residue windows (Figure 2E-F), where each window corresponds to the mutational sensitivity of approx. three turns of the helix. The relative sensitivity of each 11-residue window was remarkably consistent across concentration regimes, indicating conservation of the amphiphilic helix structure and dynamics. We therefore do not see any evidence in this model for change in the binding pose of α-synuclein as a function of concentration. Moreover, we conclude that the increasing dynamics toward the C terminus of the amphiphilic helix, which manifest as decreasing mutational sensitivity, do not result from self-interactions but rather appear intrinsic to this protein structure. Increased dynamics toward the C terminus of the helix might result from the inability of the helix to fully bury hydrophobic side chains as they wrap around the surface of the helix, due to mismatch between the periodicity of the hydrophobic groups (3.67 residues) and that of the α-helix (3.6) residues (Bussell Jr. et al., 2005; Alderson & Markley, 2013; Newberry et al., 2020).

### Changes in protein concentration shift the conformational equilibrium of specific variants

Having established that the membrane-bound helix is the dominant toxic conformation of α-synuclein at varying expression levels in yeast, we next sought to identify how changes in concentration alter the cellular population of the membrane-bound helix and its equilibrium with any coexisting structures (e.g., the unbound, disordered monomer).

At concentrations far from the dissociation constant, changes in concentration alter the total population without substantially changing the ratio of bound to unbound states, whereas at concentrations close the dissociation constant, changes in concentration cause significant changes in the bound fraction of the protein, shifting equilibrium between the bound and unbound states. If changes in concentration shift the equilibrium between bound and unbound states, then mutations that reverse this shift should exhibit concentration dependence. For example, whereas at high expression levels we were only able to discover mutations that disrupt membrane binding and toxicity, a shift toward the unbound state at lower expression levels should reveal mutations that enhance membrane binding and therefore toxicity. This scenario in particular would provide new insights into the sequence determinants of α-synuclein–membrane interactions.

In contrast, even at the lowest expression levels, we failed to identify any mutations that substantially enhance toxicity via membrane binding; instead, at all concentrations tested, we only identify mutations that disrupt toxicity. We therefore conclude that at the expression levels accessed by our experiments, the protein concentration remains significantly above the dissociation constant for membrane binding, creating a significant bias toward the membrane-bound state, which can be disrupted but not significantly enhanced. Consistent with this conclusion, previous imaging studies have shown that the subcellular localization of α-synuclein in yeast is highly biased toward cellular membranes (Outeiro & Lindquist, 2003; Soper et al., 2008).

Though changes in concentration did not convert any substitution into an enhancer of toxicity, changes in concentration did affect specific sets of variants differentially, as evident from the deviations in the linear relationship between fitness scores for each variant at different expression levels (i.e., 1% [w/v] galactose vs 0.01% [w/v] galactose) (Figure 3A). To identify variants whose behavior might change as a function of concentration, we must first account for the different selective pressure of experiments performed at different inducer concentrations. To enable comparison between datasets with distinct selective pressures, we first calculate the residual in each fitness score, relative to the linear regression between all fitness scores in each condition. When the selective pressure differs, as it does here, the slope of the linear regression diverges from one, reflecting the difference in selective pressure and resulting change in dynamic range of the fitness scores. By subtracting out that linear correlation, we can look past the difference in selective pressure and more directly compare the effect of individual mutations between conditions. Larger residuals reflect variants whose behavior deviates more from the average behavior of the population as a whole.

**Figure 3.**
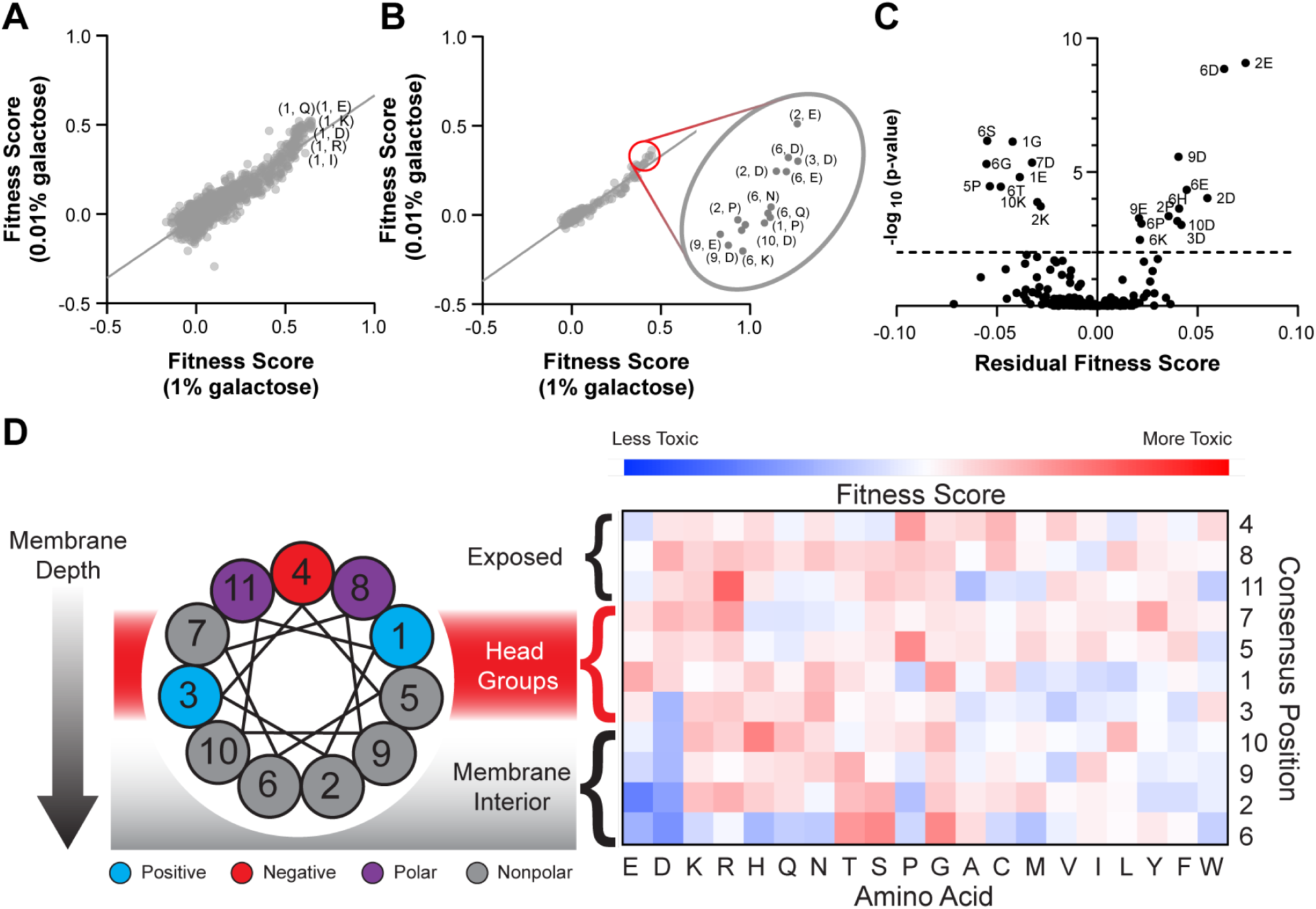
Comparison of *α*-synuclein fitness scores at high vs moderate expression levels. **(A**) Fitness scores of all variants induced with 0.01% galactose compared to the fitness scores under induction at 1% galactose. **(B)** Average fitness scores of equivalent substitutions at repeated positions in the membrane-binding region (e.g., the average effect of substituting an aspartate as the second position of each 11-residue repeat). **(C)** Volcano plot of residuals in the fitness scores plotted in panel B, relative to the linear regression between those fitness scores at different expression levels. P-values were calculated from one-sample t-test. The dashed line shows 5% false discovery rate (FDR) threshold calculated using the Benjamini-Hochberg procedure. **(D)** Residuals in the average fitness scores of equivalent substitutions at repeated positions in the membrane-binding region, relative to the linear regression between those fitness scores at different expression levels (i.e., inducer concentrations of 0.01% vs 1% galactose).

Specifically, low-toxicity variants were observed to be even more fit (i.e., less toxic) at the lower expression level than expected from the otherwise linear correlation. The same trend was observed in both the fitness of individual variants, as well as the average fitness scores of equivalent substitutions in repeated segments (e.g., the average effect of incorporating an aspartate at the sixth position of each of the seven 11-residue segments) (Figure 3A-B). To identify variants whose concentration-dependent toxicity deviates significantly from the rest of the population of variants, we calculated the residual in fitness score of each variant from the linear trend of the whole population (Figure S8, Table S3).

This analysis highlighted specific concentration-dependent effects on variants with reduced membrane affinity (Figure S9-10). For example, variants with anionic substitutions on the hydrophobic, membrane-contacting face of the helix (e.g., aspartate substitutions at the sixth position of the 11-residue repeating module), which have significantly reduced membrane affinity, were even less toxic than expected upon reducing the expression level (Figure 3C-D, Table S4). As a result of the reduced membrane affinity, these variants are likely expressed at levels closer to their dissociation constant for membrane binding, so changes in concentration are more likely to shift the equilibrium between the membrane-bound helix and unbound, disordered state, which would explain the distinct concentration dependence of reduced-affinity variants relative to variants with high membrane affinity. This shift in the equilibrium between membrane-bound and unbound states due to mutations that alter membrane affinity is also consistent with the greater cytosolic localization of α-synuclein variants with reduced membrane affinity, as determined in previous imaging experiments (Outeiro & Lindquist, 2003; Newberry et al., 2020).

Similarly, variants with substitutions of the initiator methionine, which are already expressed at much lower levels than other variants (Newberry et al., 2020a), are perturbed more strongly than variants with WT-like expression (Figure 3A). The low intrinsic expression of these variants likely brings their cellular concentration closer to the range of the dissociation constant for membrane binding, such that changes in concentration due to inducer titration are sufficient to shift the population away from the membrane-bound state, which is the main source of α-synuclein toxicity in this system. We therefore conclude that what unites the variants with the greatest concentration-dependent changes in toxicity is that they are expressed at levels closer to their respective dissociation constants for membrane binding (Figure S11).

Interestingly, the variants most differentially perturbed by changes in expression level are precisely those whose toxicity is selectively perturbed by rapamycin treatment (Figure S12). We had previously treated yeast cells with a variety of tool compounds, including rapamycin, in order to alter the folding environment within yeast cells and reveal alternative toxic states (Newberry et al., 2020a). Rapamycin treatment alters a variety of fundamental homeostatic processes, including protein and nucleic acid synthesis, autophagy, and cell cycle progression (Li et al., 2014). We previously suggested that the effects of rapamycin on the relative toxicity of α-synuclein variants in yeast was dominated by its effects on protein synthesis and therefore the steady-state expression level of α-synuclein.

The remarkable similarity in the effects of rapamycin treatment and reduced expression on the relative toxicity of α-synuclein variants further corroborates our conclusion that the effects of rapamycin on α-synuclein toxicity in yeast are dominated by rapamycin’s combined effects on protein synthesis and degradation.

### Concentration-dependent reversal in the relative effects of mutations that reduce membrane binding

At the lowest inducer concentration, there is minimal selective pressure, which allows us to sample the full range of expression levels in this cellular model. Upon reducing the inducer concentration from 1% to 0.01% (w/v) galactose, low-toxicity variants exhibited a larger change in toxicity than variants with WT-like toxicity (see above and Figure 3). Interestingly, at 0.001% (w/v) galactose, low-toxicity variants were *more* toxic than expected from the overall change in fitness of the entire population (Figure 4). A scatterplot relating the fitness score of each variant expressed at an inducer concentration of 0.001% (w/v) galactose vs at 0.01% (w/v) galactose showed significant deviation from linearity (Figure 4A-B), with low-toxicity variants being less fit (i.e., more toxic) at the lower expression level than expected from the otherwise linear correlation.

**Figure 4.**
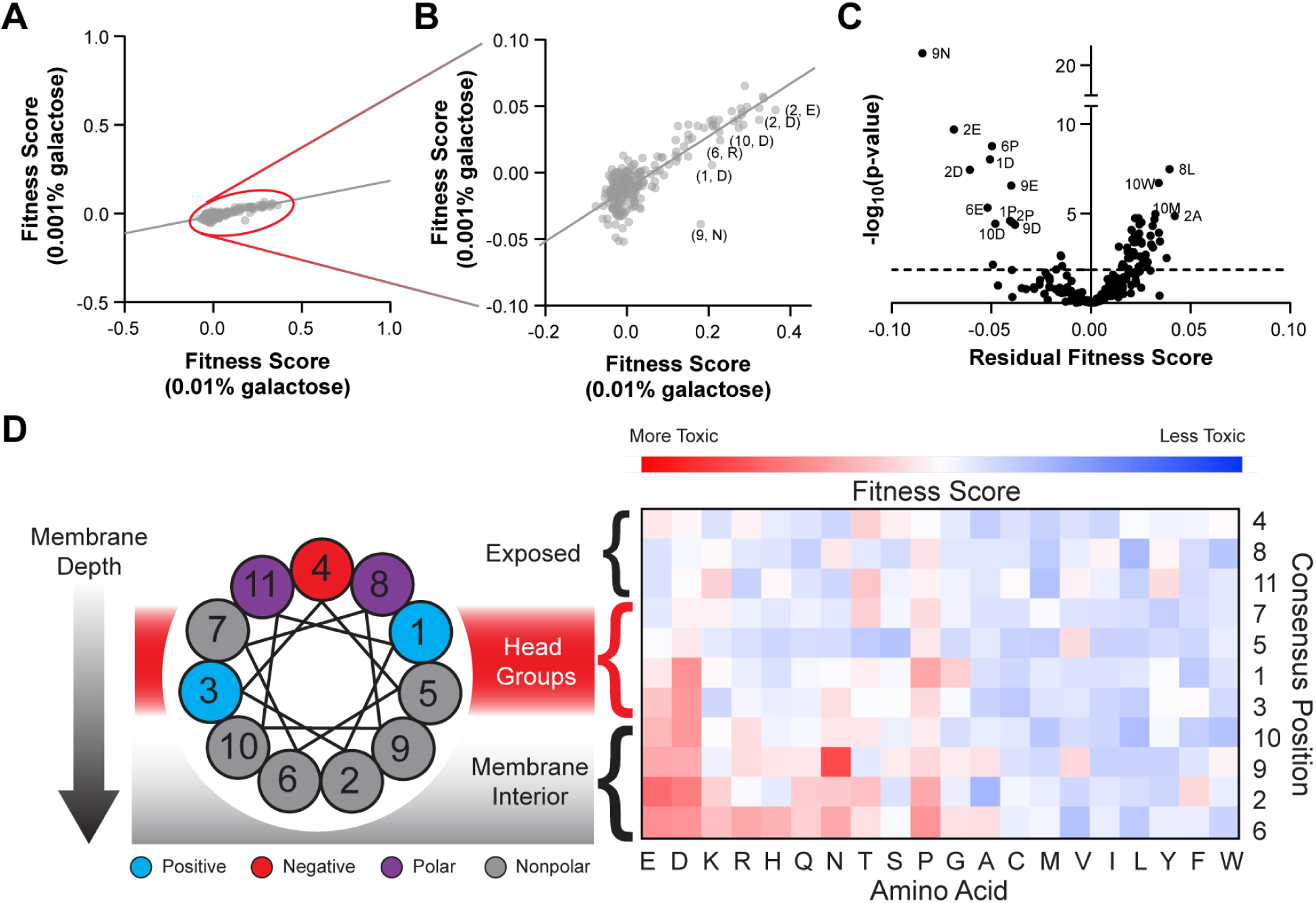
Analysis of fitness scores of *α*-synuclein variants between 0.001% and 0.01% galactose. **(A-B)** Average fitness scores of equivalent substitutions at repeated positions in the membrane-binding region (e.g., the average effect of substituting an aspartate as the second position of each 11-residue repeat) induced with 0.01% galactose compared to the fitness scores under induction at 0.001% galactose. **(C)** Volcano plot of residuals in the fitness scores plotted in panel B, relative to the linear regression between those fitness scores at different expression levels. P-values were calculated from one-sample t-test. The dashed line shows 5% false discovery rate (FDR) threshold calculated using the Benjamini-Hochberg procedure. **(D)** Residuals in the average fitness scores of equivalent substitutions at repeated positions in the membrane-binding region, relative to the linear regression between those fitness scores at different expression levels (i.e., inducer concentrations of 0.001% vs 0.01% galactose).

Upon calculating the residual in fitness score from the linear relationship for each variant or set of variants (e.g., variants with aspartate substitutions at the sixth position of each 11-residue membrane-binding segment; Figure 4C-D), we observed that the differentially perturbed variants are likely to have reduced membrane affinity due to incorporation of an anionic residue on the hydrophobic face of the amphiphilic helix. The same variants that were *less* toxic than expected upon decreasing from high to moderate expression are therefore *more* toxic than expected upon decreasing from moderate to low expression. Though the fitness scores at low expression levels are relatively small in magnitude, confidence is increased by integrating the fitness scores of variants with distinct barcodes mapping to identical amino-acid substitutions (see Methods for more details). Moreover, we further integrated fitness scores of variants with substitutions at equivalent positions within α-synuclein’s repeating sequence. As a result, each fitness score in Figure 4C/D is the composite of generally over 100 measurements each, which allows us to improve signal to noise and confidently identify variants with significantly perturbed fitness.

We offer two possible explanations for the observation that low-toxicity variants were more toxic than expected based on the change in toxicity of the overall population upon reduction to low concentration. First, it is possible that at low expression and with mutations that disrupt membrane binding, there is insufficient population of the membrane-bound state of these variants to impart significant toxicity. Whereas one would expect a reduction in concentration to reduce population of the toxic species, a low-affinity variant might not populate the toxic state substantially, so a reduction in concentration would not result in the expected change in toxicity. Alternatively, we speculate that this result would be consistent with a change in the cooperativity of membrane binding upon mutation. If binding to the membrane is negatively cooperative, then the binding of variants with lower membrane affinity might be disproportionately affected due to insufficient release of free energy upon binding to overcome repulsive interactions in the membrane-bound state. Reducing the expression level would relieve the negative cooperativity, which could disproportionately affect variants with low membrane affinity, which might otherwise be unable to engage in membrane binding and therefore toxicity. Ultimately, these data are only suggestive of cooperative interactions, which would require additional future investigation. Interestingly, previous *in vitro* studies have suggested the possibility of positive cooperativity in the binding of α-synuclein to model membranes (Makasewicz et al., 2021), which appears inconsistent with our results, suggesting different behaviors *in vitro* and in cells.

## Discussion

By comprehensively mapping the sequence determinants of functional protein folding *in situ*, deep mutational scanning offers unique insights into the conformational landscape of proteins in living cells. Given the known concentration dependence of α-synuclein folding/misfolding, we applied DMS to probe concentration-dependent changes in the cellular folding landscape, which is difficult to assess by other methods. Our results show that the membrane-bound helix is the dominant toxic conformation of α-synuclein in yeast, as changes in concentration did not reveal any gross or subtle changes in conformation, as evidenced in the concentration-dependent fitness scores. Even at low expression levels, we did not identify any substitutions that significantly increase toxicity, indicating that α-synuclein is highly biased toward the toxic, membrane-bound state in yeast. Membrane-binding capacity has also been implicated as a key driver of toxicity in other α-synuclein species, such as soluble oligomers and amyloid fibers, including those derived from phase-separated condensates (Fusco et al., 2017; Chen et al., 2024). We also identified concentration-dependent changes in the population of the toxic state that were specific to variants with reduced membrane affinity, which further emphasized the robust formation of the toxic membrane-bound helix by WT α-synuclein upon heterologous expression in yeast.

This study is also one of relatively few DMS studies performed at distinct protein expression levels. One provocative example was the recent DMS of ubonodin, an antimicrobial peptide, which leveraged different expression levels to identify variants with increased potency (Thokkadam et al., 2023). Our study highlights the further potential of concentration-dependent DMS to probe changes in the population of different conformational states in living cells, which we expect will be valuable in the study of a wide variety of targets, especially for proteins prone to aggregation. Studies like this are especially important given that different cell types can harbor dramatically different α-synuclein expression levels, which could result in distinct folding landscapes. For example, microglia and brain macrophages express much lower α-synuclein than neurons and oligodendrocytes (Wilhelm et al., 2014; Zhang et al., 2016). Given the known concentration-dependence of α-synuclein pathology in humans (Singleton et al., 2003; Ibáñez et al., 2004; Ikeuchi et al., 2008), we anticipate that studies like this one will give insight into disease-associated changes in α-synuclein folding.

## Materials and methods

### Yeast Growth Assay

*S. cerevisiae* W303 strains expressing either EGFP or α-synuclein-EGFP from a 2μ plasmid under control of the GAL promoter were inoculated into SCR-Ura (to select for transformed cells using the *URA3* prototrophy marker) and maintained in log phase overnight at 30 °C with 225 rpm shaking. Cells were then diluted into fresh SCR-Ura containing galactose at concentration from 0 to 1% (w/v). Growth of the cells was then monitored by recording the OD600 every 30 minutes using a Biotek Synergy H1 plate reader. Experiments were performed in triplicate.

### Fluorescence microscopy

*S. cerevisiae* W303 strains expressing either EGFP or α-synuclein-EGFP were inoculated into SCR-Ura and maintained in log phase overnight at 30 °C with 225 rpm shaking. Cells were then diluted into fresh SCR-Ura containing galactose at concentration from 0 to 1% (w/v). After 6 hours, the cells were imaged on an EVOS FL Auto 2 microscope equipped with a Plan fluor LWD EVOS 60x/0.75 air objective. Both brightfield and fluorescent (482 excitation and 524 emission) images were acquired.

### Library construction and cloning

Library construction was performed as described previously (Newberry et al., 2020; Newberry et al., 2020a). Briefly, a linear, double-stranded oligonucleotide library expressing all possible single-amino acid substitutions was synthesized by Twist Bioscience. This fragment was cloned into a yeast 2μ plasmid upstream of GFP and a random nucleotide barcode using the EZClone method. Restrictive transformation of the products into *E. coli* resulted in approximately 60,000 clones, and subsequent barcode association by long-read sequencing resulted in 31,484 barcode–variant mappings covering 2,600 α-synuclein variants, corresponding to approximately 12 independent barcodes per variant. The resulting plasmid library was then transformed into *S. cerevisiae* strain W303 to generate the yeast α-synuclein library.

### α-Synuclein Variant −Barcode Association

Barcode sequences uniquely coupled to individual α-synuclein variants were previously established (Newberry et al., 2020). Plasmids recovered from DH5α cells were PCR-amplified using primers targeting the α-synuclein−GFP−barcode region and incorporating Illumina adapter sequences. The resulting amplicons were gel-purified and subjected to sequencing on an Illumina MiSeq platform with custom primers for Read1, Index Read, and Read 2, run for 305, 20, and 205 cycles, respectively. Read 1 and Read 2 covered the α-synuclein coding region in forward and reverse orientations, whereas the Index Read resolved the barcode sequences. Sequencing clusters whose barcode calls carried >5% estimated probability of containing at least one error as inferred from quality scores were excluded for further analysis. Paired-end reads were merged to reconstruct full-length α-synuclein sequences. Base calls for nonoverlapping regions were taken directly from the corresponding reads, while overlapping segments were resolved by consensus. In case of disagreement, the WT base was retained when present in one of the reads. Otherwise, the base with the higher quality score was selected. Merged sequences were aligned to the designed library using Bowtie2 to generate a list of barcode-variant associations on a per-read basis. For each barcode, all observed variants were collected across the dataset. Barcode-variant associations were considered high confidence only when the most frequent assignment occurred at least two- fold more often than the second most frequent. This procedure yielded a final dictionary linking 31,484 unique barcodes to 2,600 α-synuclein variants.

### Yeast Library Selection Experiments

Outgrowth selection of the yeast library of α-synuclein variants was performed as described previously (Newberry et al., 2020). Briefly, a glycerol stock of the yeast α-synuclein library (described above) was thawed into SCD-Ura and incubated overnight at 30 °C, 225 rpm. Cells were then back diluted into SCD-Ura for an additional 12 hours of recovery. Cells were then washed and resuspended in SCR-Ura and maintained in log phase. Before induction, an aliquot of the unselected library was collected by centrifugation and stored at –80 °C. Log-phase cells were then diluted into SCR-Ura containing galactose at one of four different concentrations (i.e., 1%, 0.1%, 0.01%, and 0.001% [w/v]). After 12 hours, another aliquot of cells was collected, and the culture was diluted to maintain log phase growth. A final aliquot was collected 12 hours later. Selection experiments were performed in duplicate starting from new glycerol stocks. Plasmid DNA was then isolated from frozen cell pellets by miniprep (Fowler et al., 2014).

### Deep Sequencing

PCR was used to amplify DNA barcodes from the plasmid DNA isolated from each aliquot of the yeast library during outgrowth selection (see above). A nested PCR strategy appended the adapters and primer binding sites necessary for Illumina sequencing (Table S5). Products were gel extracted, quantified, pooled, and finally sequenced by NovaSeq in SR100 mode.

### Calculation of Fitness Scores

Fitness scores were calculated as described previously (Newberry et al., 2020). Barcode sequences with high-confident assignments to specific protein variants were quantified at each sampling time (i.e., before selection, after the first 12 hours, and after 24 hours of selection). Barcodes with fewer than 10 counts in the unselected sample were excluded from analysis. For normalization, a single representative WT barcode was chosen. This barcode was identified by computing the distribution of changes in log-transformed frequency over time for all WT-associated barcodes and selecting the one whose temporal trajectory correspond to the median of this distribution. This median WT barcode served as the reference for all subsequent comparison. At each time point, counts for barcodes linked to mutant variants were normalized to the count of the median WT barcode. The resulting mutant-to-WT ratios were then log-transformed to account for the exponential growth dynamics of yeast cells. For each barcode, the log-transformed frequencies across the three time points were fitted using a linear model, and the resulting slopes were averaged across experimental replicates. Sloes corresponding to synonymous barcodes (i.e., independent barcodes mapping to the same amino-acid variant) were further averaged to yield a fitness score and associated error for each α-synuclein variants. Statistical significance was assessed by performing Welch’s t-tests comparing fitness score of each variant to that of WT.

### Residual Analysis for Changes in Fitness Score Caused by Galactose Concentration

Fitness scores from parallel experiments were compared as previously described (Newberry et al., 2020a). To quantify the linear relationship between fitness measurements obtained under two galactose concentrations (e.g., 1% vs 0.01% [w/v] gal), we employed orthogonal distance regression, which accounts for uncertainty in both variables. Regression parameters (slope and intercept) were used to calculate an “expected” fitness score under reduced galactose conditions, reflecting the fitness predicted from the altered growth rates. For each mutant, deviations from this predicted linear relationship (i.e., residuals) were calculated under the galactose concentration (Figure 3A). These residuals represent the changes in each mutant’s fitness score attributable to perturbations in expression levels. The statistical significance of the residual fitness scores for each variant was assessed using Welch’s t-tests comparing its fitness score to that of WT.

To evaluate positional trends across the 11-residue repeats (e.g., incorporating an aspartate residue at the sixth position of each membrane-binding repeat), we averaged the residual fitness scores of equivalent substitutions (Figures 3D and 4D). Associated errors were propagated from the variances of underlying fitness measurements. The significance of these averaged positional effects was determined using one-sample t-tests on the residual fitness scores, yielding the p-values reported in Figures 3C and 4C. A 5% false discovery rate (FDR) threshold was applied by determining the p-value cutoff at which 5% of significant hits would be expected to represent false positives under the reduced protein expression conditions, as determined using the Benjamini–Hochberg procedure.

## Data availability

Raw sequencing data are available at the NCBI Sequence Read Archive (PRJNA1310013, PRJNA564806). Barcode counts in each condition at each time point for each replicate are available as supporting tables, as are the resulting fitness scores. The residual fitness scores with associated statistics are available in the supporting tables. Fitness scores are also available at MaveDB (mavedb.org; accession numbers mavedb:00001249-a-1 through mavedb:00001249-a-4). The remaining data underlying this study are available in the article and its supporting information.

## Code availability

Programs developed to analyze the data reported are available at github.com/rnewberry17.

## Supplementary Material Description

**Figure S1.** Growth of W303 strains expressing either EGFP or α-synuclein-EGFP at different galactose concentrations.

**Figure S2.** Fluorescence micrographs of W303 cells expressing α-synuclein-EGFP under selected galactose concentrations.

**Figure S3.** Correlation of fitness scores of all α-synuclein variants between duplicate selection experiments conducted under varying galactose concentrations.

**Figure S4.** Growth of W303 strains expressing selected variants at different galactose concentrations.

**Figure S5.** Average fitness scores of variants with hydrophobic, polar, or proline across 140 residues of α-synuclein under four different expression levels.

**Figure S6.** Fitness scores of known familial Parkinson’s disease variants.

**Figure S7.** Average fitness scores of all 19 substitutions at each position of α-synuclein, calculated for selection experiments at each of four different protein expression levels.

**Figure S8.** Residuals in the fitness scores of α-synuclein variants compared to the linear regression of fitness scores between different expression levels. Four pairwise comparisons are presented.

**Figure S9.** Sensitivity to changes in galactose concentrations.

**Figure S10.** Residuals in the average fitness scores of equivalent substitutions at repeated positions in the membrane-binding region.

**Figure S11.** Cellular toxicity is dependent on the membrane-bound population of α-synuclein.

**Figure S12.** Fitness scores and volcano plot of α-synuclein variants in rapamycin treatment.

**Figure S13.** Distribution of mutant fitness scores as a function of inducer concentration.

**Table S1.** Counts of each variant at each time point

**Table S2.** Table of complete fitness scores

**Table S3.** Table of complete residual fitness scores

**Table S4.** Table of residual fitness scores and statistical significance

**Table S5.** List of DNA Sequences

**Table S6.** Fitness scores of familial Parkinson’s disease variants at different protein expression levels

## Acknowledgements

This work was supported by grants from the NIH (R00-NS116679 and R35-GM160477) to RWN. We thank staff from the UT Austin Genomic Sequencing and Analysis Facility (GSAF) for support in next-generation sequencing.

## References

1. Afitska K, Fucikova A, Shvadchak VV, Yushchenko DA. α-Synuclein aggregation at low concentrations. Biochim Biophys Acta Proteins Proteom. 2019; 1867(7): 701–709.

2. Alderson TR, Markley JL. Biophysical characterization of α-synuclein and its controversial structure. Intrinsically Disord Proteins. 2013; 1(1): e26255.

3. Arutyunyan A, Seuma M, Faure AJ, Bolognesi B, Lehner B. Massively parallel genetic perturbation suggests the energetic structure of an amyloid-β transition state. Sci Adv. 2025; 11(24): eadv1422.

4. Bendor JT, Logan TP, Edwards RH. The function of α-synuclein. Neuron. 2013; 79(6): 1044–1066.

5. Binolfi A, Limatola A, Verzini S, Kosten J, Theillet FX, Rose HM, Bekei B, Stuiver M, van Rossum M, Selenko P. Intracellular repair of oxidation-damaged α-synuclein fails to target C-terminal modification sites. Nat Commun. 2016; 7(1): 10251.

6. Bodner CR, Dobson CM, Bax A. Multiple tight phospholipid-binding modes of α-synuclein revealed by solution NMR spectroscopy. J Mol Biol. 2009; 390(4): 775–790.

7. Bolognesi B, Faure AJ, Seuma M, Schmiedel JM, Tartaglia GG, Lehner B. The mutational landscape of a prion-like domain. Nat Commun. 2019; 10(1): 4162.

8. Bradbury J. α-Synuclein gene triplication discovered in Parkinson’s disease. Lancet Neurol. 2003; 2(12): 715.

9. Buell AK, Galvagnion C, Gaspar R, Sparr E, Vendruscolo M, Knowles TPJ, Linse S, Dobson CM. Solution conditions determine the relative importance of nucleation and growth processes in α-synuclein aggregation. Proc Natl Acad Sci USA. 2014; 111(21): 7671–7676.

10. Burré J, Sharma M, Südhof TC. α-Synuclein assembles into higher-order multimers upon membrane binding to promote SNARE complex formation. Proc Natl Acad Sci USA. 2014; 111(40): E4274–E4283.

11. Busch DJ, Oliphint PA, Walsh RB, Banks SML, Woods WS, George JM, Morgan JR. Acute increase of α-synuclein inhibits synaptic vesicle recycling evoked during intense stimulation. Mol Biol Cell. 2014; 25(24): 3926–3941.

12. Bussell R Jr, Ramlall TF, Eliezer D. Helix periodicity, topology, and dynamics of membrane-associated α-synuclein. Protein Sci. 2005; 14(4): 862–872.

13. Bussell R, Eliezer D. A structural and functional role for 11-mer repeats in α-synuclein and other exchangeable lipid binding proteins. J Mol Biol. 2003; 329(4): 763–778.

14. Calabresi P, Mechelli A, Natale G, Volpicelli-Daley L, Di Lazzaro G, Ghiglieri V. α- Synuclein in Parkinson’s disease and other synucleinopathies: from overt neurodegeneration back to early synaptic dysfunction. Cell Death Dis. 2023; 14(3): 176.

15. Chen SW, Barritt JD, Cascella R, Bigi A, Cecchi C, Banchelli M, Gallo A, Jarvis JA, Chiti F, Dobson CM, Fusco G, De Simone A. Structure–toxicity relationship in intermediate fibrils from α-synuclein condensates. J Am Chem Soc. 2024;146(15): 10537–10549.

16. Chiti F, Dobson CM. Protein misfolding, amyloid formation, and human disease: a summary of progress over the last decade. Annu Rev Biochem. 2017; 86: 27–68.

17. Chlebowicz J, Russ W, Chen D, Vega A, Vernino S, White III CL, Rizo J, Joachimiak LA, Diamond MI. Saturation mutagenesis of α-synuclein reveals monomer fold that modulates aggregation. Sci Adv. 2023; 9(43): eadh3457.

18. Chung CY, Khurana V, Auluck PK, Tardiff DF, Mazzulli JR, Soldner F, Baru V, Lou Y, Freyzon Y, Cho S, et al. Identification and rescue of α-synuclein toxicity in Parkinson patient-derived neurons. Science. 2013; 342(6161): 983–987.

19. Conway KA, Harper JD, Lansbury PT Jr. Accelerated in vitro fibril formation by a mutant α-synuclein linked to early-onset Parkinson disease. Nat Med. 1998; 4(11): 1318–1320.

20. Cooper AA, Gitler AD, Cashikar A, Haynes CM, Hill KJ, Bhullar B, Liu K, Xu K, Strathearn KE, Liu F, et al. α-Synuclein blocks ER-Golgi traffic and Rab1 rescues neuron loss in Parkinson’s models. Science. 2006; 313(5785): 324–328.

21. Davidson WS, Jonas A, Clayton DF, George JM. Stabilization of α-synuclein secondary structure upon binding to synthetic membranes. J Biol Chem. 1998; 273(16): 9443–9449.

22. Delenclos M, Burgess JD, Lamprokostopoulou A, Outeiro TF, Vekrellis K, McLean PJ. Cellular models of α-synuclein toxicity and aggregation. J Neurochem. 2019; 150(5): 566–576.

23. Dettmer U, Newman AJ, von Saucken VE, Bartels T, Selkoe D. KTKEGV repeat motifs are key mediators of normal α-synuclein tetramerization: their mutation causes excess monomers and neurotoxicity. Proc Natl Acad Sci USA. 2015; 112(31): 9596–9601.

24. Dettmer U, Ramalingam N, von Saucken VE, Kim TE, Newman AJ, Terry-Kantor E, Selkoe D. Loss of native α-synuclein multimerization by strategically mutating its amphipathic helix causes abnormal vesicle interactions in neuronal cells. Hum Mol Genet. 2017; 26(18): 3466–3481.

25. Di Maio R, Barrett PJ, Hoffman EK, Barrett CW, Zharikov A, Borah A, Hu X, McCoy J, Chu CT, Burton EA, et al. α-Synuclein binds to TOM20 and inhibits mitochondrial protein import in Parkinson’s disease. Sci Transl Med. 2016; 8(342): 342ra78.

26. Dumarieh R, Lagasca D, Krishna S, Kragelj J, Xiao Y, Ansari S, Frederick KK. Structural context modulates the conformational ensemble of the intrinsically disordered amino terminus of α-synuclein. J Am Chem Soc. 2025; 147(14): 11800–11810.

27. Eliezer D, Kutluay E, Bussell R, Browne G. Conformational properties of α-synuclein in its free and lipid-associated states. J Mol Biol. 2001; 307(4): 1061–1073.

28. Fanning S, Haque A, Imberdis T, Baru V, Barrasa MI, Nuber S, Termine D, Ramalingam N, Ho GPH, Noble T, et al. Lipidomic analysis of α-synuclein neurotoxicity identifies stearoyl CoA desaturase as a target for Parkinson treatment. Mol Cell. 2019; 73(5): 1001–1014.e8.

29. Fares MB, Jagannath S, Lashuel HA. Reverse engineering Lewy bodies: how far have we come and how far can we go? Nat Rev Neurosci. 2021; 22(2): 111–131.

30. Fowler DM, Fields S. Deep mutational scanning: a new style of protein science. Nat Methods. 2014; 11(8): 801–807.

31. Fowler DM, Stephany JJ, Fields S. Measuring the activity of protein variants on a large scale using deep mutational scanning. Nat Protoc. 2014; 9(9): 2267–2284.

32. Fusco G, Chen SW, Williamson PTF, Cascella R, Perni M, Jarvis JA, Cecchi C, Vendruscolo M, Chiti F, Cremades N, et al. Structural basis of membrane disruption and cellular toxicity by α-synuclein oligomers. Science. 2017; 358(6369): 1440–1443.

33. Fusco G, De Simone A, Gopinath T, Vostrikov V, Vendruscolo M, Dobson CM, Veglia G. Direct observation of the three regions in α-synuclein that determine its membrane-bound behaviour. Nat Commun. 2014; 5(1): 3827.

34. Fusco G, Pape T, Stephens AD, Mahou P, Costa AR, Kaminski CF, Kaminski Schierle GS, Vendruscolo M, Veglia G, Dobson CM, et al. Structural basis of synaptic vesicle assembly promoted by α-synuclein. Nat Commun. 2016; 7(1): 12563.

35. Gallardo R, Ranson NA, Radford SE. Amyloid structures: much more than just a cross-β fold. Curr Opin Struct Biol. 2020; 60: 7–16.

36. Galvagnion C, Buell AK, Meisl G, Michaels TCT, Vendruscolo M, Knowles TPJ, Dobson CM. Lipid vesicles trigger α-synuclein aggregation by stimulating primary nucleation. Nat Chem Biol. 2015; 11(3): 229–234.

37. Georgieva ER, Ramlall TF, Borbat PP, Freed JH, Eliezer D. Membrane-bound α-synuclein forms an extended helix: long-distance pulsed ESR measurements using vesicles, bicelles, and rodlike micelles. J Am Chem Soc. 2008; 130(39): 12856–12857.

38. Gitler AD, Bevis BJ, Shorter J, Strathearn KE, Hamamichi S, Su LJ, Caldwell KA, Caldwell GA, Rochet JC, McCaffery JM, et al. The Parkinson’s disease protein α-synuclein disrupts cellular Rab homeostasis. Proc Natl Acad Sci USA. 2008; 105(1): 145–150.

39. Gitler AD, Chesi A, Geddie ML, Strathearn KE, Hamamichi S, Hill KJ, Caldwell KA, Caldwell GA, Cooper AA, Rochet JC, et al. α-Synuclein is part of a diverse and highly conserved interaction network that includes PARK9 and manganese toxicity. Nat Genet. 2009; 41(3): 308–315.

40. Gray VE, Sitko K, Kameni FZN, Williamson M, Stephany JJ, Hasle N, Fowler DM. Elucidating the molecular determinants of Aβ aggregation with deep mutational scanning. G3 (Bethesda). 2019; 9(11): 3683–3689.

41. Ibáñez P, Bonnet AM, Débarges B, Lohmann E, Tison F, Agid Y, Dürr A, Brice A, Pollak P; French Parkinson’s Disease Genetics Study Group. Causal relation between α-synuclein locus duplication as a cause of familial Parkinson’s disease. Lancet. 2004; 364(9440): 1169–1171.

42. Ikeuchi T, Kakita A, Shiga A, Kasuga K, Kaneko H, Tan CF, Idezuka J, Wakabayashi K, Onodera O, Iwatsubo T, et al. Patients homozygous and heterozygous for SNCA duplication in a family with parkinsonism and dementia. Arch Neurol. 2008; 65(4): 514–519.

43. Iljina M, Garcia GA, Horrocks MH, Klenerman D, et al. Kinetic model of the aggregation of α-synuclein provides insights into prion-like spreading. Proc Natl Acad Sci USA. 2016; 113(9): E1206–E1215.

44. Jao CC, Der-Sarkissian A, Chen J, Langen R. Structure of membrane-bound α-synuclein studied by site-directed spin labeling. Proc Natl Acad Sci USA. 2004; 101(22): 8331–8336.

45. Jao CC, Hegde BG, Chen J, Haworth IS, Langen R. Structure of membrane-bound α-synuclein from site-directed spin labeling and computational refinement. Proc Natl Acad Sci USA. 2008; 105(50): 19666–19671.

46. Khurana V, Lindquist S. Modelling neurodegeneration in *Saccharomyces cerevisiae*: why cook with baker’s yeast? Nat Rev Neurosci. 2010; 11(6): 436–449.

47. Khurana V, Peng J, Chung CY, Auluck PK, Fanning S, Tardiff DF, Bartels T, Koeva M, Eichhorn SW, Benyamini H, et al. Genome-scale networks link neurodegenerative disease genes to α-synuclein through specific molecular pathways. Cell Syst. 2017; 4(2): 157–170.e14.

48. Kim TE, Newman AJ, Imberdis T, Brontesi L, Tripathi A, Ramalingam N, Fanning S, Selkoe D, Dettmer U. Excess membrane binding of monomeric alpha-, beta- and gamma-synuclein is invariably associated with inclusion formation and toxicity. Hum Mol Genet. 2021; 30(23): 2332–2346.

49. Knowles TPJ, Vendruscolo M, Dobson CM. The amyloid state and its association with protein misfolding diseases. Nat Rev Mol Cell Biol. 2014; 15(6): 384–396.

50. Konno T, Ross OA, Puschmann A, Dickson DW, Wszolek ZK. Autosomal dominant Parkinson’s disease caused by SNCA duplications. Parkinsonism Relat Disord. 2016; 22(Suppl 1): S1–S6.

51. Kragelj J, Ghosh R, Xiao Y, Dumarieh R, Lagasca D, Krishna S, Frederick KK. Spatially resolved DNP-assisted NMR illuminates the conformational ensemble of α-synuclein in intact viable cells. Proc Natl Acad Sci USA. 2025; 122(23): e2500367122.

52. Lashuel HA. Membrane permeabilization: a common mechanism in protein-misfolding diseases. Sci Aging Knowl Environ. 2005; 2005(38): p e28.

53. Lashuel HA, Overk CR, Oueslati A, Masliah E. The many faces of α-synuclein: from structure and toxicity to therapeutic target. Nat Rev Neurosci. 2013; 14(1): 38–48.

54. Li D, Liu C. Conformational strains of pathogenic amyloid proteins in neurodegenerative diseases. Nat Rev Neurosci. 2022; 23(9): 523–534.

55. Li J, Kim SG, Blenis J. Rapamycin: one drug, many effects. Cell Metab. 2014; 19(3): 373–379.

56. Louros N, Schymkowitz J, Rousseau F. Mechanisms and pathology of protein misfolding and aggregation. Nat Rev Mol Cell Biol. 2023; 24(12): 912–933.

57. Luk KC, Kehm V, Carroll J, Zhang B, O’Brien P, Trojanowski JQ, Lee VM. Pathological α-synuclein transmission initiates Parkinson-like neurodegeneration in non-transgenic mice. Science. 2012; 338(6109): 949–953.

58. Makasewicz K, Wennmalm S, Stenqvist B, Fornasier M, Andersson A, Jönsson P, Linse S, Sparr E. Cooperativity of α-Synuclein Binding to Lipid Membranes. ACS Chem. Neurosci. 2021; 12(12): 2099–2109

59. Makasewicz K, Carlström G, Stenström O, Bernfur K, Fridolf S, Akke M, Linse S, Sparr E. Tipping point in α-synuclein-membrane interactions from stable protein-covered vesicles to amyloid aggregation. Cell Rep. Phys. Sci. 2024; 5(12): 102309.

60. Mansueto S, Fusco G, De Simone A. α-Synuclein and biological membranes: the danger of loving too much. Chem Commun (Camb). 2023; 59(57): 8769–8778.

61. Meisl G, Kirkegaard JB, Arosio P, Michaels TCT, Vendruscolo M, Dobson CM, Linse S, Knowles TPJ. Molecular mechanisms of protein aggregation from global fitting of kinetic models. Nat Protoc. 2016; 11(2): 252–272.

62. Milchberg MH, Warmuth OA, Borcik CG, Dhavale DD, Wright ER, Kotzbauer PT, Rienstra CM. α-Synuclein fibril structures cluster into distinct classes. Biophys J. 2025; 124(16): 2571–2582.

63. Narhi L, Wood SJ, Steavenson S, Jiang Y, Wu GM, Anafi D, et al. Both familial Parkinson’s disease mutations accelerate α-synuclein aggregation. J Biol Chem. 1999; 274(14): 9843–9846.

64. Newberry RW, Leong JT, Chow ED, Kampmann M, DeGrado WF. Deep mutational scanning reveals the structural basis for α-synuclein activity. Nat Chem Biol. 2020; 16(6): 653–659.

65. Newberry RW, Arhar T, Costello J, Hartoularos GC, Maxwell AM, Naing ZZC, Pittman M, Reddy NR, Schwarz DM, Wassarman DR, et al. Robust sequence determinants of α-synuclein toxicity in yeast implicate membrane binding. ACS Chem Biol. 2020a; 15(8): 2137–2153.

66. Outeiro T, Lindquist S. Yeast cells provide insight into α-synuclein biology and pathobiology. Science. 2003; 302(5651): 1772-5.

67. Perrino G, Wilson C, Santorelli M, Di Bernardo D. Quantitative characterization of α-synuclein aggregation in living cells through automated microfluidics feedback control. Cell Rep. 2019; 27(3): 916–927.e5.

68. Roeters SJ, Strunge K, Pedersen KB, Golbek TW, Bregnhøj M, Zhang Y, Wang Y, Dong M, Nielsen J, Otzen DE, et al. Elevated concentrations cause upright α-synuclein conformation at lipid interfaces. Nat Commun. 2023; 14(1): 5731.

69. Rollins NJ, Brock KP, Poelwijk FJ, Stiffler MA, Gauthier NP, Sander C, Marks DS. Inferring protein 3D structure from deep mutation scans. Nat Genet. 2019; 51(7): 1170–1176.

70. Sampaio-Marques B, Guedes A, Vasilevskiy I, Gonçalves S, Outeiro TF, Winderickx J, Burhans WC, Ludovico P. α-Synuclein toxicity in yeast and human cells is caused by cell cycle re-entry and autophagy degradation of ribonucleotide reductase 1. Aging Cell. 2019; 18(4): e12922.

71. Sangkaew A, Kojornna T, Tanahashi R, Takagi H, Yompakdee C. A novel yeast-based screening system for potential compounds that can alleviate human α-synuclein toxicity. J Appl Microbiol. 2022; 132(2): 1409–1421.

72. Schmiedel JM, Lehner B. Determining protein structures using deep mutagenesis. Nat Genet. 2019; 51(7): 1177–1186.

73. Seuma M, Lehner B, Bolognesi B. An atlas of amyloid aggregation: the impact of substitutions, insertions, deletions and truncations on amyloid beta fibril nucleation. Nat Commun. 2022; 13(1): 7084.

74. Shahmoradian SH, Lewis AJ, Genoud C, Hench J, Moors TE, Navarro PP, Castaño-Díez D, Schweighauser G, Graff-Meyer A, Goldie KN, et al. Lewy pathology in Parkinson’s disease consists of crowded organelles and lipid membranes. Nat Neurosci. 2019; 22(7): 1099–1109.

75. Singleton AB, Farrer M, Johnson J, Singleton A, Hague S, Kachergus J, Hulihan M, Peuralinna T, Dutra A, Nussbaum R, et al. α-Synuclein locus triplication causes Parkinson’s disease. Science. 2003; 302(5646): 841.

76. Snead D, Eliezer D. Intrinsically disordered proteins in synaptic vesicle trafficking and release. J Biol Chem. 2019; 294(10): 3325–3342.

77. So RWL, Watts JC. α-Synuclein conformational strains as drivers of phenotypic heterogeneity in neurodegenerative diseases. J Mol Biol. 2023; 435(12): 168011.

78. Soper JH, Roy S, Stieber A, Lee E, Wilson RB, Trojanowski JQ, Burd CG, Lee VM. α-Synuclein–induced aggregation of cytoplasmic vesicles in *Saccharomyces cerevisiae*. Mol Biol Cell. 2008; 19(3): 1093–1103.

79. Theillet FX, Binolfi A, Bekei B, Martorana A, Rose HM, Stuiver M, Verzini S, Lorenz D, van Rossum M, Goldfarb D, et al. Structural disorder of monomeric α-synuclein persists in mammalian cells. Nature. 2016; 530(7588): 45–50.

80. Thokkadam A, Do T, Ran X, Brynildsen MP, Yang ZJ, Link AJ. High-throughput screen reveals the structure–activity relationship of the antimicrobial lasso peptide ubonodin. ACS Cent Sci. 2023; 9(3): 540–550.

81. Thompson M, Martín M, Olmo TS, Rajesh C, Koo PK, Bolognesi B, Lehner B. Massive experimental quantification allows interpretable deep learning of protein aggregation. Sci Adv. 2025; 11(18): eadt5111.

82. Tosatto L, Andrighetti AO, Plotegher N, Antonini V, Tessari I, Ricci L, Bubacco L, Dalla Serra M. α-Synuclein pore forming activity upon membrane association. Biochim Biophys Acta Biomembr. 2012; 1818(11): 2876–2883.

83. Trinkaus VA, Riera-Tur I, Martínez-Sánchez A, Bäuerlein FJB, Guo Q, Arzberger T, Baumeister W, Dudanova I, Hipp MS, Hartl FU, et al. In situ architecture of neuronal α-synuclein inclusions. Nat Commun. 2021; 12(1): 2110.

84. Tsuboyama K, Dauparas J, Chen J, Laine E, Mohseni Behbahani Y, Weinstein JJ, Mangan NM, Ovchinnikov S, Rocklin GJ. Mega-scale experimental analysis of protein folding stability in biology and design. Nature. 2023; 620(7973): 434–444.

85. Vamvaca K, Volles MJ, Lansbury PT Jr. The first N-terminal amino acids of α-synuclein are essential for α-helical structure formation in vitro and membrane binding in yeast. J Mol Biol. 2009; 389(2): 413–424.

86. Villar-Piqué A, Lopes da Fonseca T, Outeiro TF. Structure, function and toxicity of α-synuclein: the Bermuda triangle in synucleinopathies. J Neurochem. 2015; 133(5): 584–589.

87. Wei H, Li X. Deep mutational scanning: a versatile tool in systematically mapping genotypes to phenotypes. Front Genet. 2023; 14: 1087267.

88. Weinreb PH, Zhen W, Poon AW, Conway KA, Lansbury PT. NACP, a protein implicated in Alzheimer’s disease and learning, is natively unfolded. Biochemistry. 1996; 35(43): 13709–13715.

89. Wilhelm BG, Mandad S, Truckenbrodt S, Kröhnert K, Schäfer C, Rammner B, Koo SJ, Claßen GA, Krauss M, Haucke V, et al. Composition of isolated synaptic boutons reveals the amounts of vesicle trafficking proteins. Science. 2014; 344(6187): 1023–1028.

90. Wong YC, Krainc D. α-Synuclein toxicity in neurodegeneration: mechanism and therapeutic strategies. Nat Med. 2017; 23(1): 1–13.

91. Wood SJ, Wypych J, Steavenson S, Louis JC, Citron M, Biere AL. α-Synuclein fibrillogenesis is nucleation-dependent: implications for the pathogenesis of Parkinson’s disease. J Biol Chem. 1999; 274(28): 19509–19512.

92. Yeger-Lotem E, Riva L, Su LJ, Gitler AD, Cashikar AG, King OD, Auluck PK, Geddie ML, Valastyan JS, Karger DR, et al. Bridging high-throughput genetic and transcriptional data reveals cellular responses to α-synuclein toxicity. Nat Genet. 2009; 41(3): 316–323.

93. Zabrocki P, Bastiaens I, Delay C, Bammens T, Ghillebert R, Pellens K, De Virgilio C, Van Leuven F, Winderickx J. Phosphorylation, lipid raft interaction and traffic of α-synuclein in a yeast model for Parkinson. Biochim Biophys Acta Mol Cell Res. 2008; 1783(10): 1767–1780.

94. Zhang Y, Sloan SA, Clarke LE, Caneda C, Plaza CA, Blumenthal PD, Vogel H, Steinberg GK, Edwards MSB, Li G, et al. Purification and characterization of progenitor and mature human astrocytes reveals transcriptional and functional differences with mouse. Neuron. 2016; 89(1): 37–53.

