## Supplementary material for "Concentration-Dependent Mutational Scanning Probes the Cellular Folding Landscape of *α*-Synuclein in Yeast": Figure S

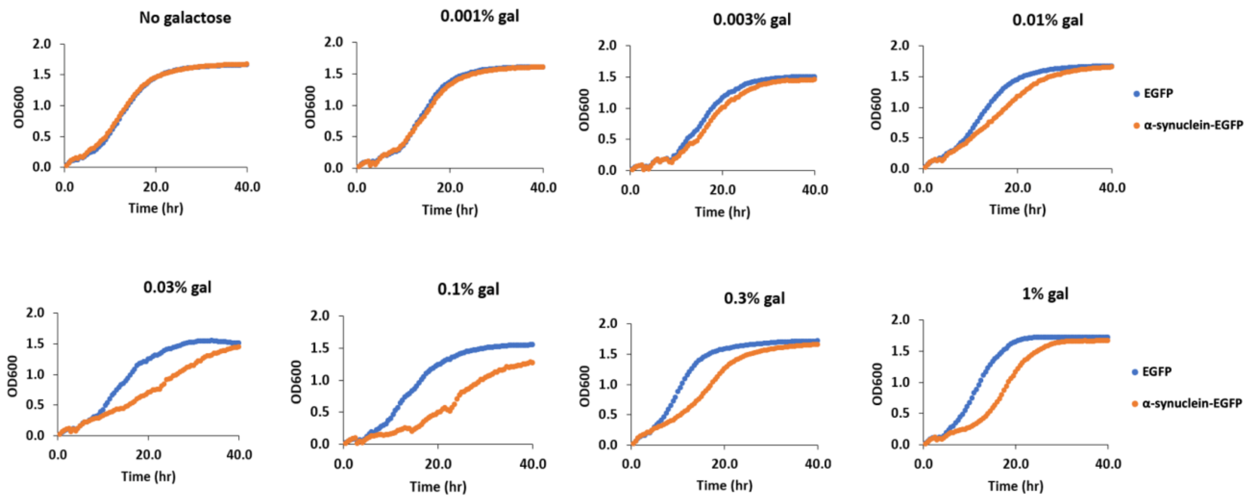


**Figure S1.** Growth of W303 strains expressing either EGFP or α-synuclein-EGFP at different galactose concentrations.

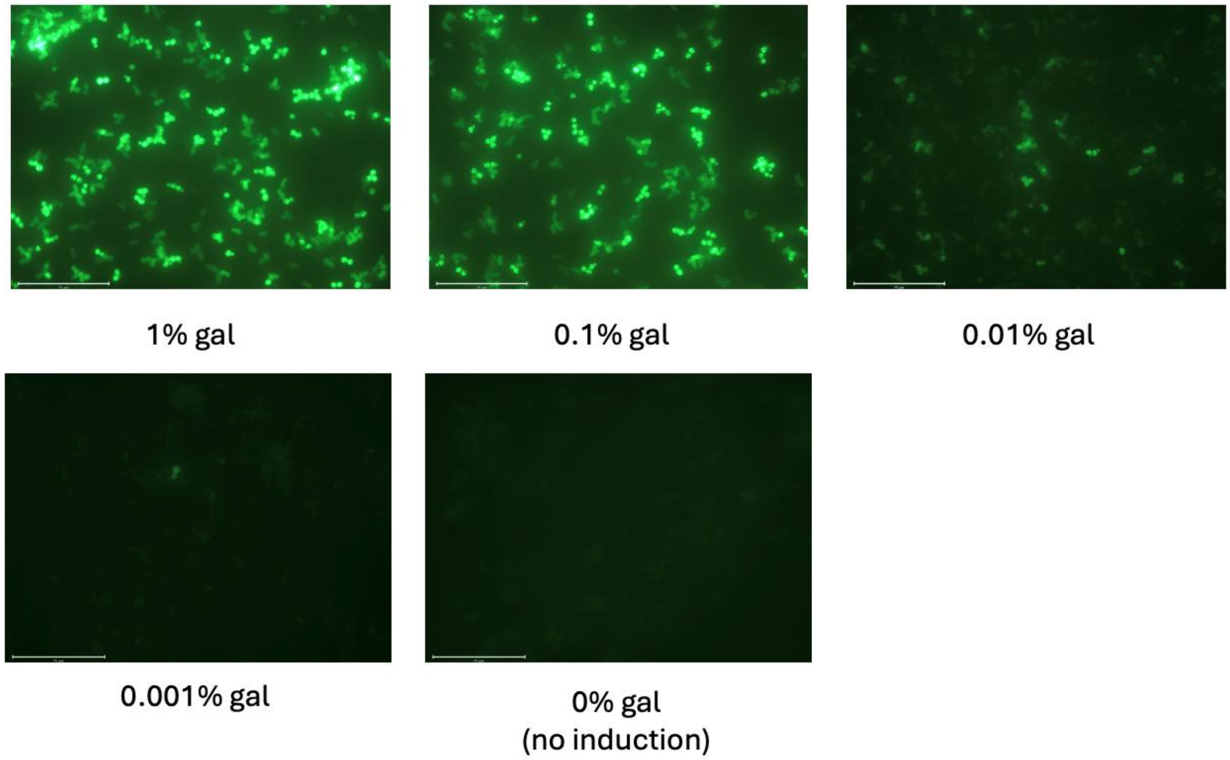


**Figure S2.** Fluorescence micrographs of W303 cells expressing α-synuclein-EGFP under selected galactose concentrations.

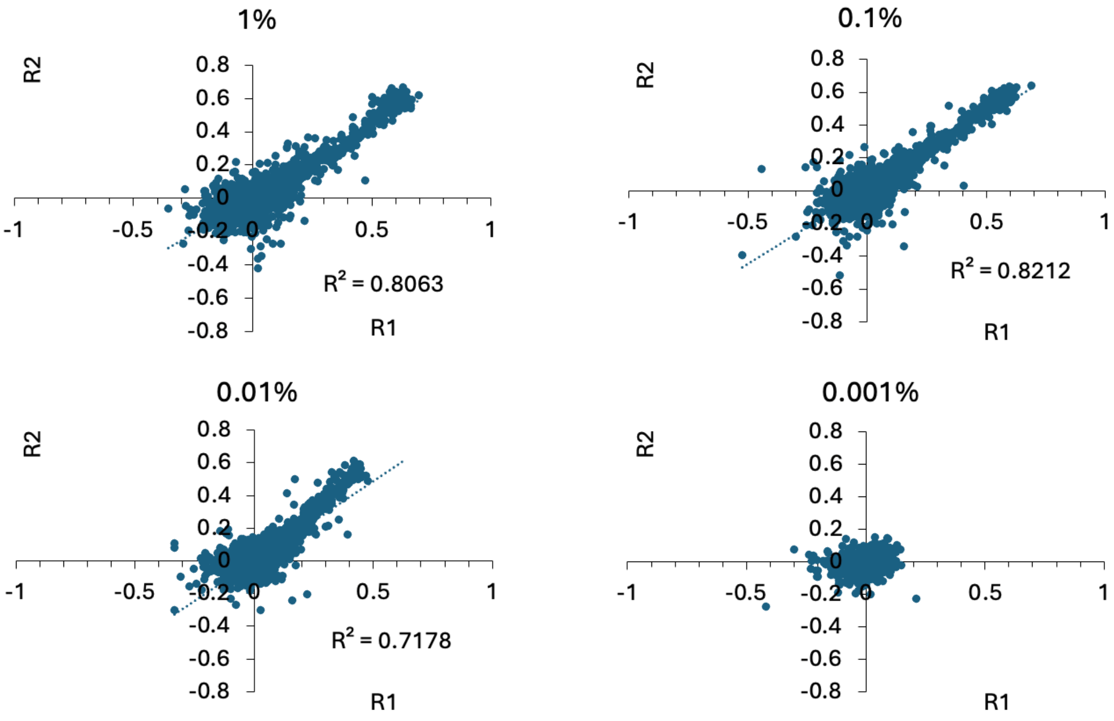


**Figure S3.** Correlation of fitness scores of all α-synuclein variants between duplicate selection experiments conducted under varying galactose concentrations.

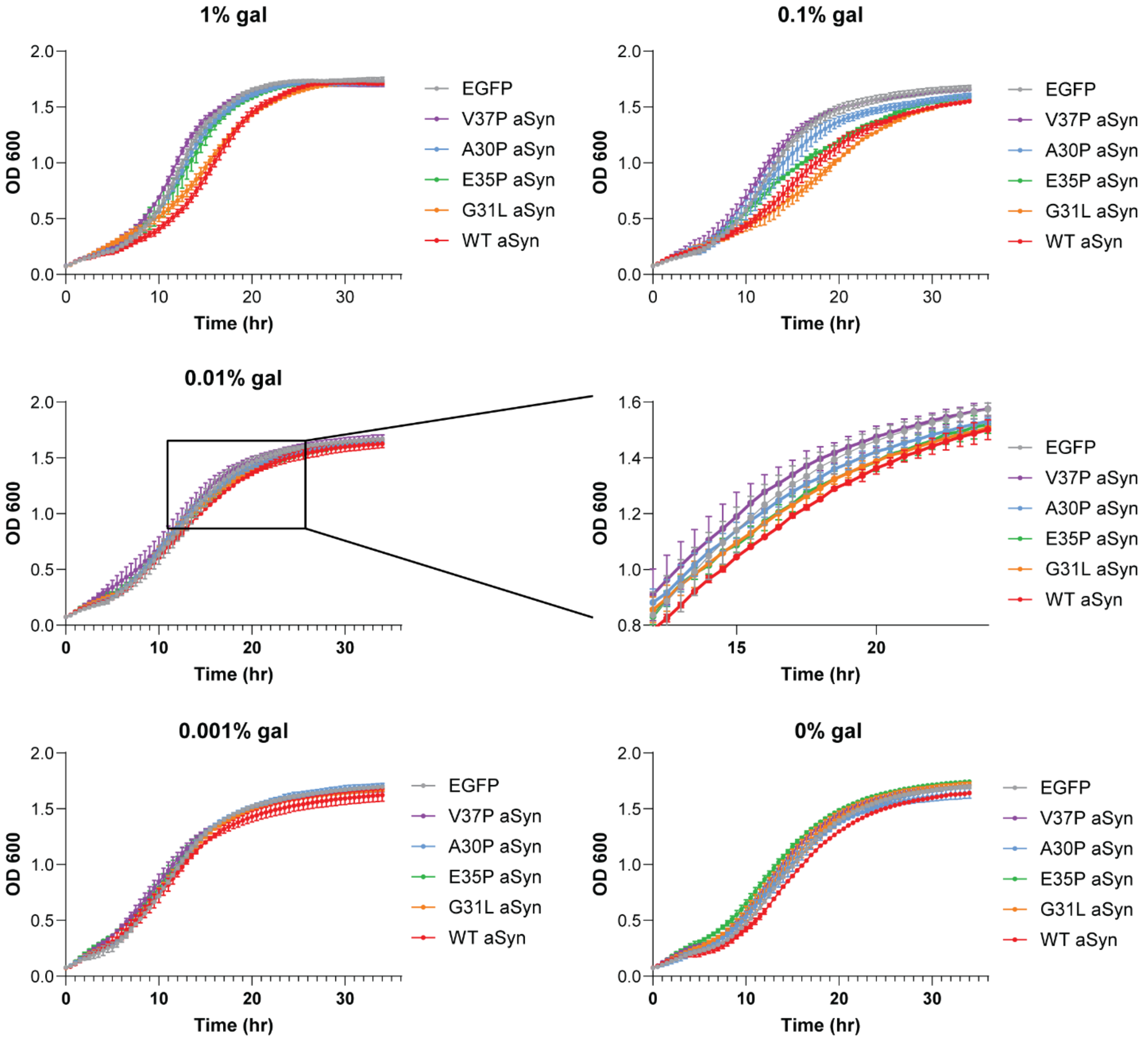


**Figure S4.** Growth of W303 strains expressing selected α-synuclein variants at different galactose concentrations.

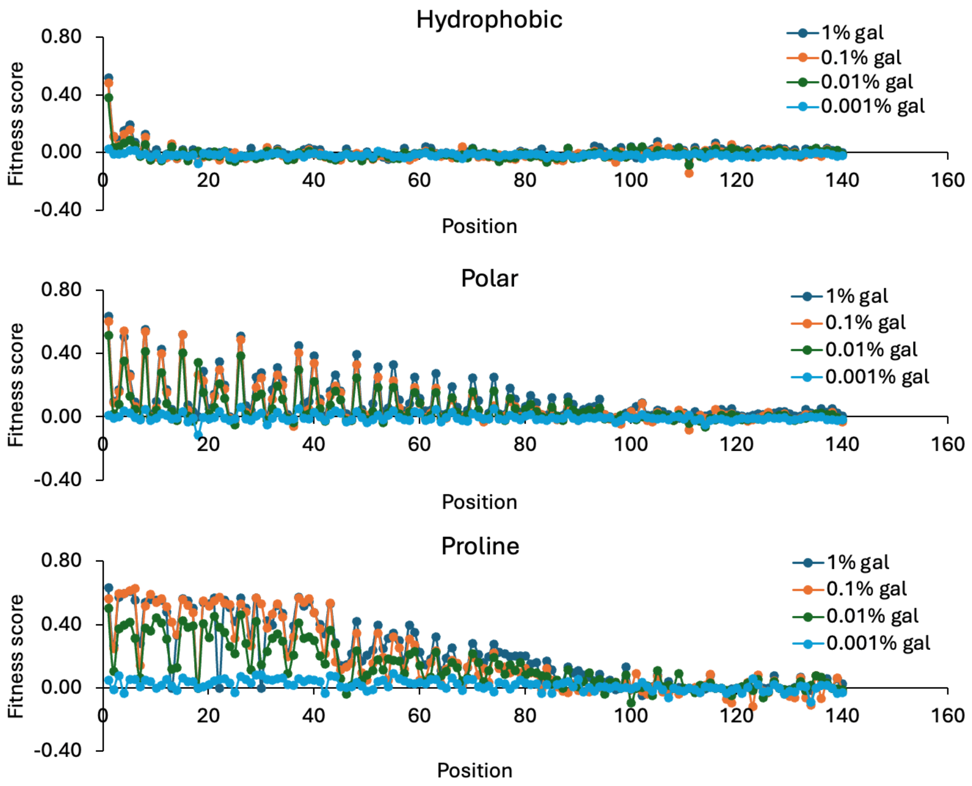


**Figure S5.** Average fitness scores of variants with hydrophobic (W, F, Y, L, I, M, C, A), polar (S, T, N, Q, H, R, K, D, E) or proline across 140 residues of α-synuclein under four different expression levels.

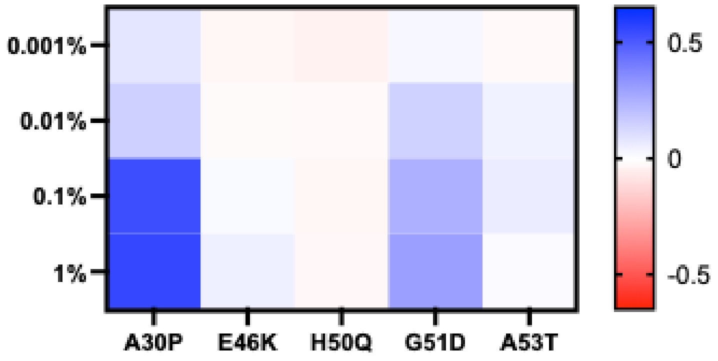


**Figure S6.** Fitness scores of known familial Parkinson’s disease variants.

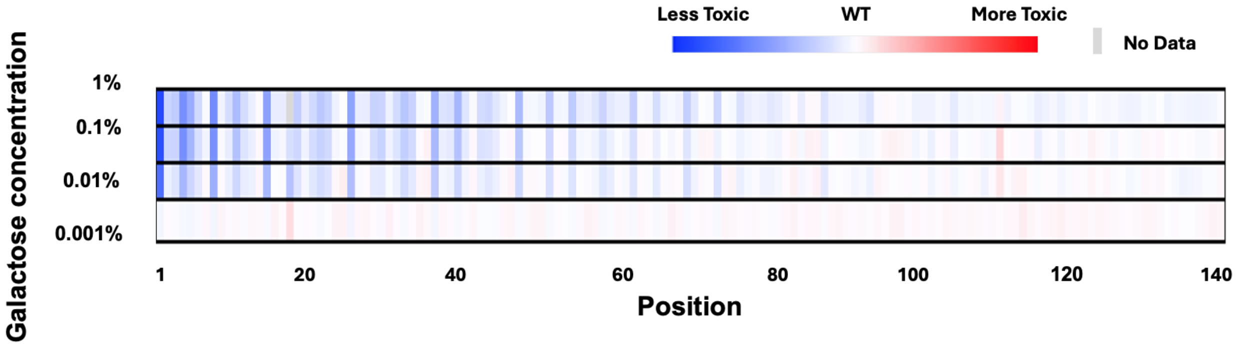


**Figure S7.** Average fitness scores of all 19 substitutions at each position of α-synuclein, calculated for selection experiments at each of four different protein expression levels.

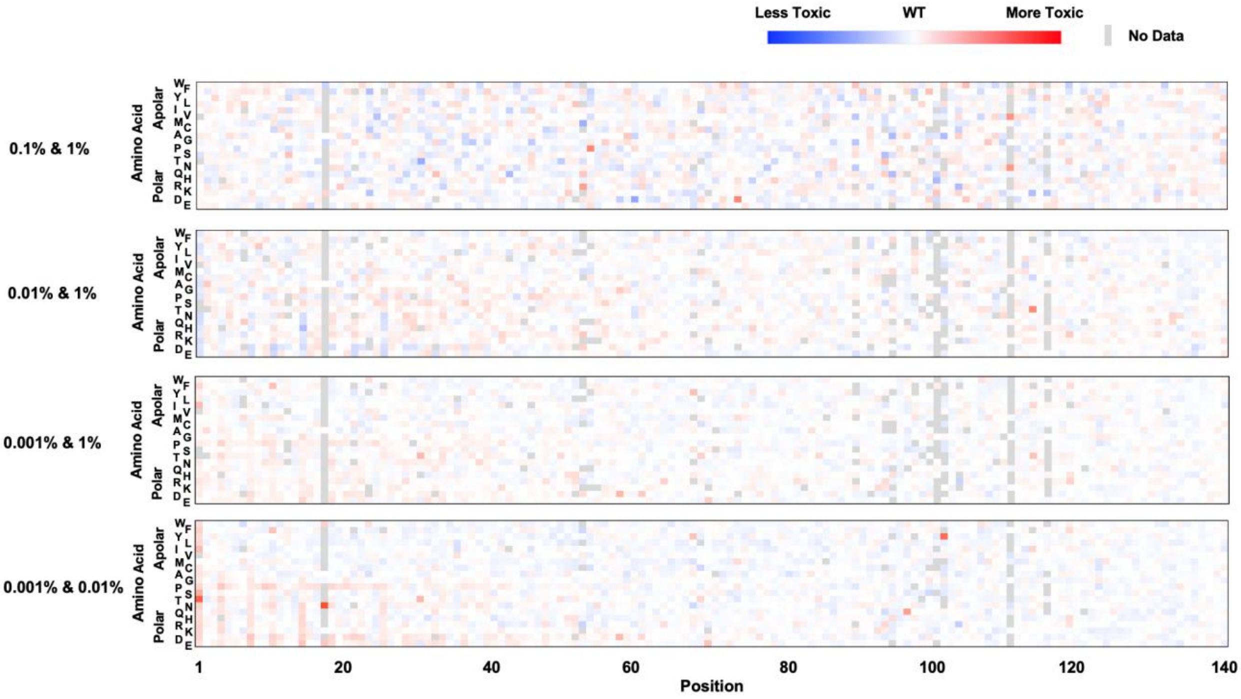


**Figure S8.** Residuals in the fitness scores of α-synuclein variants compared to the linear regression of fitness scores between different expression levels. Four pairwise comparisons are presented.

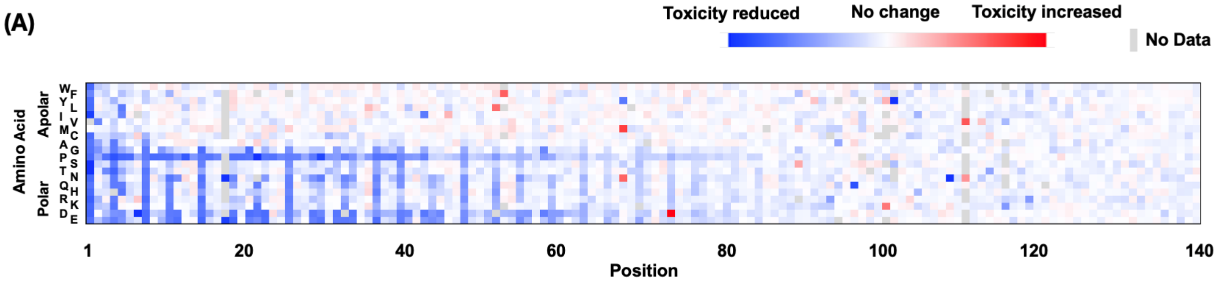


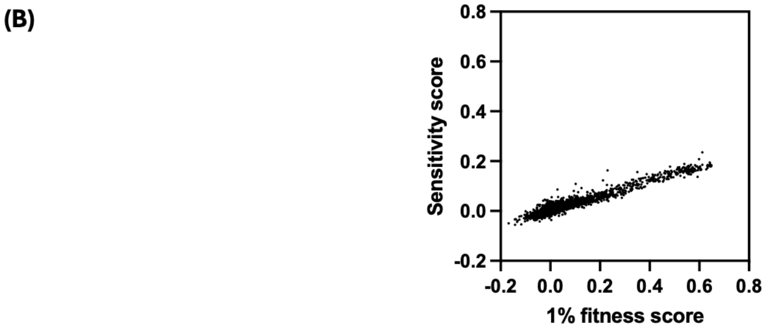


**Figure S9.** Sensitivity to changes in galactose concentrations. (A) Sensitivity scores for all α-synuclein variants. Sensitivity scores were calculated from the change in fitness score across all four inducer concentrations. Variants exhibiting increased toxicity with higher galactose concentrations are shown in blue, while those with reduced toxicity are shown in red. White indicates no significant concentration-dependent change in fitness relative to WT. (B) Correlation of sensitivity scores and fitness scores of α-synuclein variants at high expression levels.

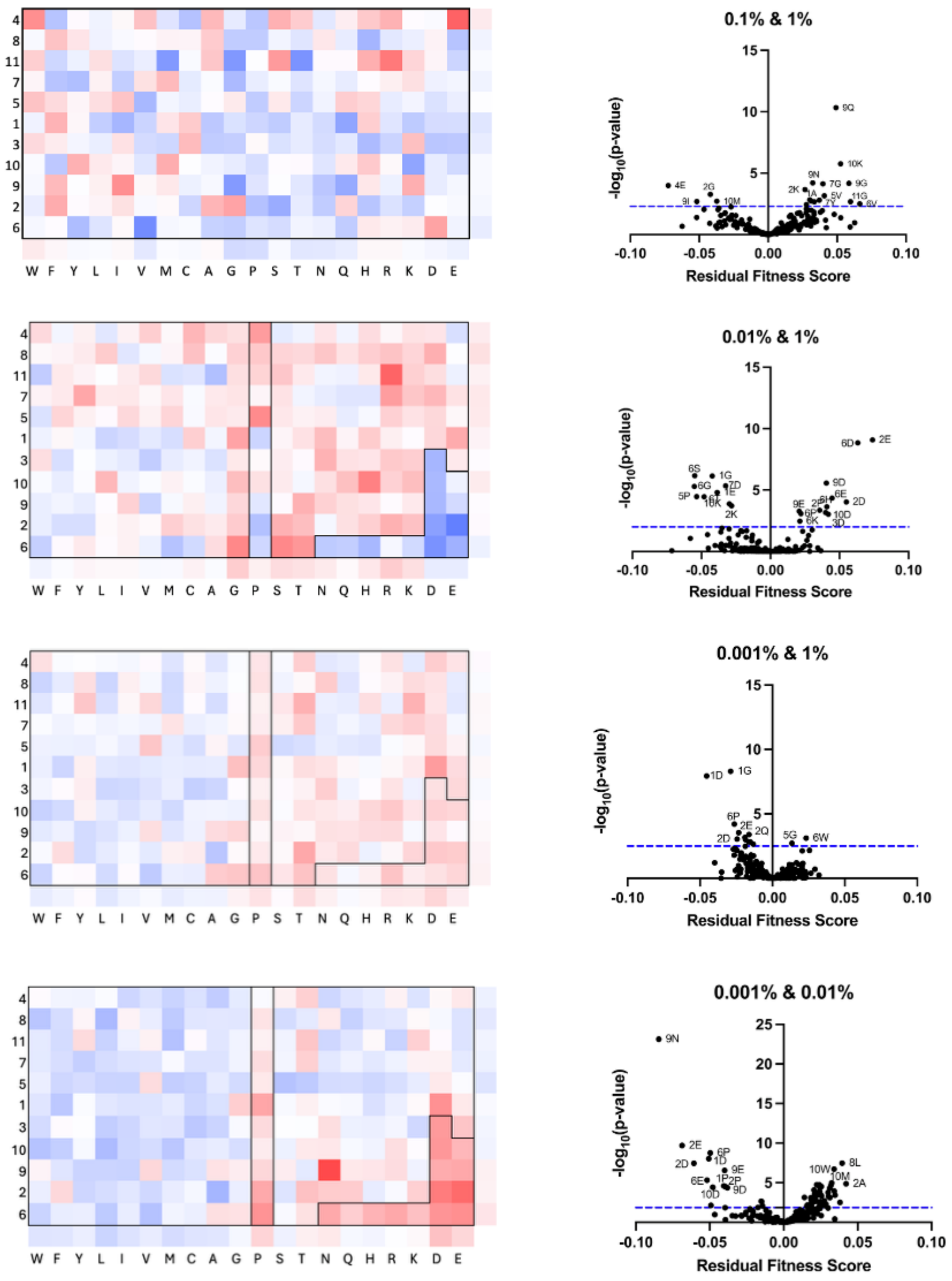


**Figure S10.** Residuals in the average fitness scores of equivalent substitutions at repeated positions in the membrane-binding region (e.g., the average effect of substituting an aspartate as the second position of each 11-residue repeat), relative to the linear regression between those fitness scores at different expression levels. The numbering of the positions is the same as in Figure 3D. A 5% false discovery rate threshold is shown as dashed blue lines in the volcano plots.

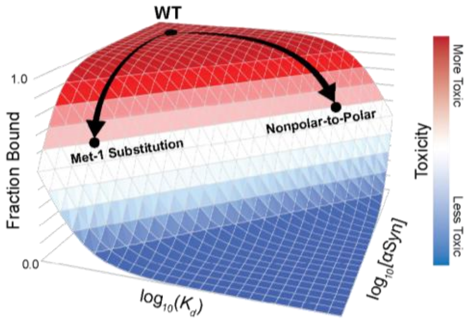


**Figure S11.** Cellular toxicity is dependent on the membrane-bound population of α-synuclein. In untreated cells, WT α-synuclein associated with cellular membranes above a critical threshold that induces toxicity. Variants can mitigate toxicity by reducing the membrane-bound fraction of α-synuclein–either by lowering membrane affinity (e.g., via the introduction of polar residues on the membrane-binding face) or by decreasing expression levels (e.g., through substitution of the initiating methionine residue, Met-1). This figure is adapted from Newberry et al. (2020), *ACS Chem Biol.*

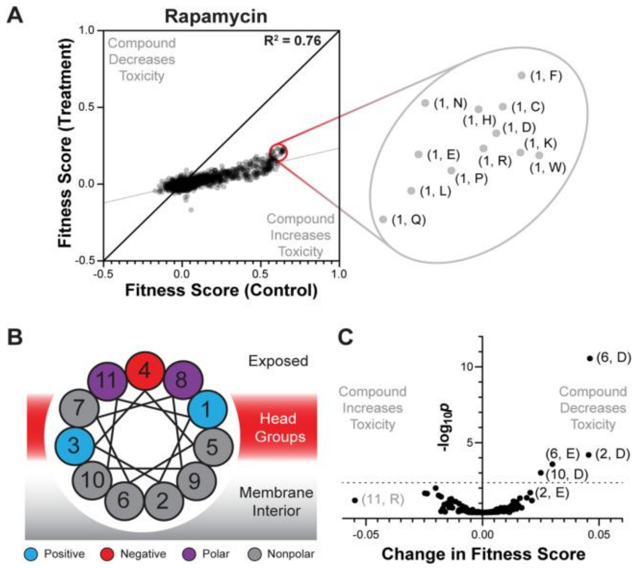


**Figure S12.** Fitness scores and volcano plot of α-synuclein variants in rapamycin treatment. (A) Fitness scores of α-synuclein variants [labeled as (position, amino-acid substitution)] in the presence of rapamycin relative to untreated controls. (B) Helical wheel of the 11-residue repeats of α-synuclein, colored by physicochemical properties. (C) Volcano plot of rapamycin-induced changes in fitness score, averaged across equivalent variants in the seven repeats denoted by (position, amino-acid substitution). This figure is adapted from Newberry et al. (2020), *ACS Chem Biol.*

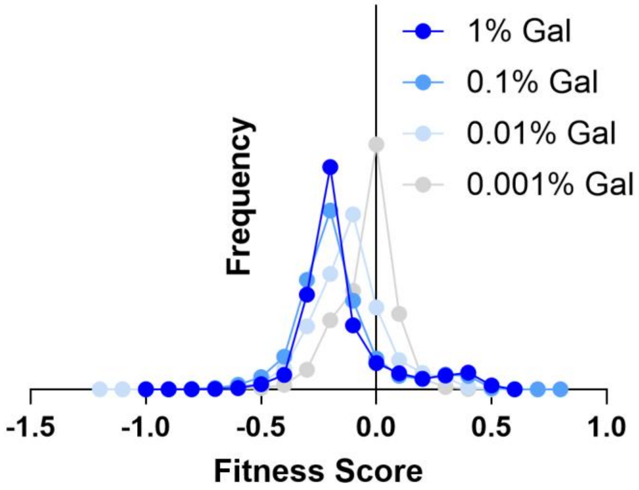


**Figure S13.** Distribution of mutant fitness scores as a function of inducer concentration.

### Table S5. List of DNA Sequences

| | |
|---|---|
| PCR 1 Forward Primer | ACACTCTTTCCCTACACGACGCTCTTCCGATCTCATGGCATGGACGAGCTGTACAAGTAA |
| PCR 1 Reverse Primer | GTGACTGGAGTTCAGACGTGTGCTCTTCCGATCTCCTTTTCGGTTAGAGCGGATGTGG |
| PCR 2 Forward Primer – UDI001 | AATGATACGGCGACCACCGAGATCTACACAGCGCTAGACACTCTTTCCCTACACGACGC |
| PCR 2 Reverse Primer – UDI001 | CAAGCAGAAGACGGCATACGAGATAACCGCGGGTGACTGGAGTTCAGACGTGTGC |
| PCR 2 Forward Primer – UDI002 | AATGATACGGCGACCACCGAGATCTACACGATATCGAACACTCTTTCCCTACACGACGC |
| PCR 2 Reverse Primer – UDI002 | CAAGCAGAAGACGGCATACGAGATGGTTATAAGTGACTGGAGTTCAGACGTGTGC |
| PCR 2 Forward Primer – UDI003 | AATGATACGGCGACCACCGAGATCTACACCGCAGACGACACTCTTTCCCTACACGACGC |
| PCR 2 Reverse Primer – UDI003 | CAAGCAGAAGACGGCATACGAGATCCAAGTCCGTGACTGGAGTTCAGACGTGTGC |
| PCR 2 Forward Primer – UDI004 | AATGATACGGCGACCACCGAGATCTACACTATGAGTAACACTCTTTCCCTACACGACGC |
| PCR 2 Reverse Primer – UDI004 | CAAGCAGAAGACGGCATACGAGATTTGGACTTGTGACTGGAGTTCAGACGTGTGC |
| PCR 2 Forward Primer – UDI005 | AATGATACGGCGACCACCGAGATCTACACAGGTGCGTACACTCTTTCCCTACACGACGC |
| PCR 2 Reverse Primer – UDI005 | CAAGCAGAAGACGGCATACGAGATCAGTGGATGTGACTGGAGTTCAGACGTGTGC |
| PCR 2 Forward Primer – UDI006 | AATGATACGGCGACCACCGAGATCTACACGAACATACACACTCTTTCCCTACACGACGC |
| PCR 2 Reverse Primer – UDI006 | CAAGCAGAAGACGGCATACGAGATTGACAAGCGTGACTGGAGTTCAGACGTGTGC |
| PCR 2 Forward Primer – UDI013 | AATGATACGGCGACCACCGAGATCTACACAAGGATGAACACTCTTTCCCTACACGACGC |
| PCR 2 Reverse Primer – UDI013 | GCACACGTCTGAACTCCAGTCACCCAAGTCTATCTCGTATGCCGTCTTCTGCTTG |
| PCR 2 Forward Primer – UDI014 | AATGATACGGCGACCACCGAGATCTACACGGAAGCAGACACTCTTTCCCTACACGACGC |
| PCR 2 Reverse Primer – UDI014 | CAAGCAGAAGACGGCATACGAGATGAGTCCAAGTGACTGGAGTTCAGACGTGTGC |
| PCR 2 Forward Primer – UDI015 | AATGATACGGCGACCACCGAGATCTACACTCGTGACCACACTCTTTCCCTACACGACGC |
| PCR 2 Reverse Primer – UDI015 | CAAGCAGAAGACGGCATACGAGATCTTAAGCCGTGACTGGAGTTCAGACGTGTGC |
| PCR 2 Forward Primer – UDI016 | ATGATACGGCGACCACCGAGATCTACACCTACAGTTACACTCTTTCCCTACACGACGC |
| PCR 2 Reverse Primer – UDI016 | CAAGCAGAAGACGGCATACGAGATTCCGGATTGTGACTGGAGTTCAGACGTGTGC |
| PCR 2 Forward Primer – UDI017 | AATGATACGGCGACCACCGAGATCTACACATATTCACACACTCTTTCCCTACACGACGC |
| PCR 2 Reverse Primer – UDI017 | CAAGCAGAAGACGGCATACGAGATCTGTATTAGTGACTGGAGTTCAGACGTGTGC |
| PCR 2 Forward Primer – UDI018 | AATGATACGGCGACCACCGAGATCTACACGCGCCTGTACACTCTTTCCCTACACGACGC |
| PCR 2 Reverse Primer – UDI018 | CAAGCAGAAGACGGCATACGAGATTCACGCCGGTGACTGGAGTTCAGACGTGTGC |
| Plasmid Sequence | CTGCATTAATGAATCGGCCAACGCGCGGGGAGAGGCGGTTTGCGTATTGGGCGCTCTTCC<br>GCTTCCTCGCTCACTGACTCGCTGCGCTCGGTCGTTCGGCTGCGGCGAGCGGTATCAGCT<br>CACTCAAAGGCGGTAATACGGTTATCCACAGAATCAGGGGATAACGCAGGAAAGAACATG<br>TGAGCAAAAGGCCAGCAAAAGCCCAGGAACCGTAAAAAGGCCGCGTTGCTGGCGTTTTTC<br>CATAGGCTCCGCCCCCCTGACGAGCATCACAAAAATCGACGCTCAAGTCAGAGGTGGCGA<br>AACCCGACAGGACTATAAAGATACCAGGCGTTTCCCCCTGGAAGCTCCCTCGTGCGCTCT<br>CCTGTTCCGACCCTGCCGCTTACCGGATACCTGTCCGCCTTTCTCCCTTCGGGAAGCGTG<br>GCGCTTTCTCATAGCTCACGCTGTAGGTATCTCAGTTCGGTGTAGGTCGTTCGCTCCAAG<br>CTGGGCTGTGTGCACGAACCCCCCGTTCAGCCCGACCGCTGCGCCTTATCCGGTAACTAT<br>CGTCTTGAGTCCAACCCGGTAAGACACGACTTATCGCCACTGGCAGCAGCCACTGGTAAC<br>AGGATTAGCAGAGCGAGGTATGTAGGCGGTGCTACAGAGTTCTTGAAGTGGTGGCCTAAC<br>TACGGCTACACTAGAAGGACAGTATTTGGTATCTGCGCTCTGCTGAAGCCAGTTACCTTC<br>GGAAAAAGAGTTGGTAGCTCTTGATCCGGCAAACAAACCACCGCTGGTAGCGGTGGTTTT<br>TTTGTTTGCAAGCAGCAGATTACGCGCAGAAAAAAAGGATCTCAAGAAGATCCTTTGATC<br>TTTTCTACGGGGTCTGACGCTCAGTGGAACGAAAACTCACGTTAAGGGATTTTGGTCATG<br>AGATTATCAAAAAGGATCTTCACCTAGATCCTTTTAAATTAAAAATGAAGTTTTAAATCA<br>ATCTAAAGTATATATGAGTAAACTTGGTCTGACAGTTACCAATGCTTAATCAGTGAGGCA<br>CCTATCTCAGCGATCTGTCTATTTCGTTCATCCATAGTTGCCTGACTCCCCGTCGTGTAG<br>ATAACTACGATACGGGAGCGCTTACCATCTGGCCCCAGTGCTGCAATGATACCGCGAGAC |

| | |
|---|---|
| | CCACGCTCACCGGCTCCAGATTTATCAGCAATAAACCAGCCAGCCGGAAGGGCCGAGCGC<br>AGAAGTGGTCCTGCAACTTTATCCGCCTCCATTCAGTCTATTAATTGTTGCCGGGAAGCT<br>AGAGTAAGTAGTTCGCCAGTTAATAGTTTGCGCAACGTTGTTGGCATTGCTACAGGCATC<br>GTGGTGTCACTCTCGTCGTTTGGTATGGCTTCATTCAGCTCCGGTTCCCAACGATCAAGG<br>CGAGTTACATGATCCCCCATGTTGTGCAAAAAAGCGGTTAGCTCCTTCGGTCCTCCGATC<br>GTTGTCAGAAGTAAGTTGGCCGCAGTGTTATCACTCATGGTTATGGCAGCACTGCATAAT<br>TCTCTTACTGTCATGCCATCCGTAAGATGCTTTTCTGTGACTGGTGAGTACTCAACCAAG<br>TCATTCTGAGAATAGTGTATGCGGCGACCGAGTTGCTCTTGCCCGGCGTCAATACGGGAT<br>AATAGTGTATCACATAGCAGAACTTTAAAAGTGCTCATCATTGGAAAACGTTCTTCGGGG<br>CGAAAACTCTCAAGGATCTTACCGCTGTTGAGATCCAGTTCGATGTAACCCACTCGTGCA<br>CCCAACTGATCTTCAGCATCTTTTACTTTCACCAGCGTTTCTGGGTGAGCAAAAACAGGA<br>AGGCAAAATGCCGCAAAAAAGGGAATAAGGGCGACACGGAAATGTTGAATACTCATACTC<br>TTCCTTTTTCAATGGGTAATAACTGATATAATTAAATTGAAGCTCTAATTTGTGAGTTTA<br>GTATACATGCATTTACTTATAATACAGTTTTTTAGTTTTGCTGGCCGCATCTTCTCAAAT<br>ATGCTTCCCAGCCTGCTTTTCTGTAACGTTCACCCTCTACCTTAGCATCCCTTCCCTTTG<br>CAAATAGTCCTCTTCCAACAATAATAATGTCAGATCCTGTAGAGACCACATCATCCACGG<br>TTCTATACTGTTGACCCAATGCGTCTCCCTTGTCATCTAAACCCACACCGGGTGTCATAA<br>TCAACCAATCGTAACCTTCATCTCTTCCACCCATGTCTCTTTGAGCAATAAAGCCGATAA<br>CAAAATCTTTGTCGCTCTTCGCAATGTCAACAGTACCCTTAGTATATTCTCCAGTAGATA<br>GGGAGCCCTTGCATGACAATTCTGCTAACATCAAAAGGCCTCTAGGTTCCTTTGTTACTT<br>CTTCTGCCGCCTGCTTCAAACCGCTAACAATACCTGGGCCCACCACACCGTGTGCATTCG<br>TAATGTCTGCCCATTCTGCTATTCTGTATACACCCGCAGAGTACTGCAATTTGACTGTAT<br>TACCAATGTCAGCAAATTTTCTGTCTTCGAAGAGTAAAAAATTGTACTTGGCGGATAATG<br>CCTTTAGCGGCTTAACTGTGCCCTCCATGGAAAAATCAGTCAAGATATCCACATGTGTTT<br>TTAGTAAACAAATTTTGGGACCTAATGCTTCAACTAACTCCAGTAATTCCTTGGTGGTAC<br>GAACATCCAATGAAGCACACAAGTTTGTTTGCTTTTCGTGCATGATATTAAATAGCTTGG<br>CAGCAACAGGACTAGGATGAGTAGCAGCACGTTCCTTATATGTAGCTTTCGACATGATTT<br>ATCTTCGTTTCCTGCAGGTTTTTGTTCTGTGCAGTTGGGTTAAGAATACTGGGCAATTTC<br>ATGTTTCTTCAACACTACATATGCGTATATATACCAATCTAAGTCTGTGCTCCTTCCTTC<br>GTTCTTCCTTCTGTTCGGAGATTACCGAATCAAAAAAATTTCAAAGAAACCGAAATCAAA<br>AAAAAGAATAAAAAAAAAAATGATGAATTGAATTGAAAAGCTAGCTTATCGATGATAAGCT<br>GTCAAAGATGAGAATTAATTCCACGGACTATAGACTATACTAGATACTCCGTCTACTGTA<br>CGATACACTTCCGCTCAGGTCCTTGTCCTTTAACGAGGCCTTACCACTCTTTTGTTACTC<br>TATTGATCCAGCTCAGCAAAGGCAGTGTGATCTAAGATTCTATCTTCGCGATGTAGTAAA<br>ACTAGCTAGACCGAGAAAGAGACTAGAAATGCAAAAGGCACTTCTACAATGGCTGCCATC<br>ATTATTATCCGATGTGACGCTGCAGCTTCTCAATGATATTCGAATACGCTTTGAGGAGAT<br>ACAGCCTAATATCCGACAAACTGTTTTACAGATTTACGATCGTACTTGTTACCCATCATT<br>GAATTTTGAACATCCGAACCTGGGAGTTTTCCCTGAAACAGATAGTATATTTGAACCTGT<br>ATAATAATATATAGTCTAGCGCTTTACGGAAGACAATGTATGTATTTCGGTTCCTGGAGA<br>AACTATTGCATCTATTGCATAGGTAATCTTGCACGTCGCATCCCCGGTTCATTTTCTGCG<br>TTTCCATCTTGCACTTCAATAGCATATCTTTGTTAACGAAGCATCTGTGCTTCATTTTGT<br>AGAACAAAAATGCAACGCGAGAGCGCTAATTTTTCAAACAAAGAATCTGAGCTGCATTTT<br>TACAGAACAGAAATGCAACGCGAAAGCGCTATTTTACCAACGAAGAATCTGTGCTTCATT<br>TTTGTAAAACAAAAATGCAACGCGACGAGAGCGCTAATTTTTCAAACAAAGAATCTGAGC<br>TGCATTTTTACAGAACAGAAATGCAACGCGAGAGCGCTATTTTACCAACAAAGAATCTAT<br>ACTTCTTTTTTGTTCTACAAAAATGCATCCCGAGAGCGCTATTTTTCTAACAAAGCATCT<br>TAGATTACTTTTTTTCTCCTTTGTGCGCTCTATAATGCAGTCTCTTGATAACTTTTTGCA<br>CTGTAGGTCCGTTAAGGTTAGAAGAAGGCTACTTTGGTGTCTATTTTCTCTTCCATAAAA<br>AAAGCCTGACTCCACTTCCCGCGTTTACTGATTACTAGCGAAGCTGCGGGTGCATTTTTT<br>CAAGATAAAGGCATCCCCGATTATATTCTATACCGATGTGGATTGCGCATACTTTGTGAA<br>CAGAAAGTGATAGCGTTGATGATTCTTCATTGGTCAGAAAATTATGAACGGTTTCTTCTA<br>TTTTGTCTCTATATACTACGTATAGGAAATGTTTACATTTTCGTATTGTTTTCGATTCAC<br>TCTATGAATAGTTCTTACTACAATTTTTTTGTCTAAAGAGTAATACTAGAGATAAACATA<br>AAAAATGTAGAGGTCGAGTTTAGATGCAAGTTCAAGGAGCGAAAGGTGGATGGGTAGGTT<br>ATATAGGGATATAGCACAGAGATATATAGCAAAGAGATACTTTTGAGCAATGTTTGTGGA<br>AGCGGTATTCGCAATGGGAAGCTCCACCCCGGTTGATAATCAGAAAAGCCCCAAAAACAG<br>GAAGATTGTATAAGCAAATATTTAAATTGTAAACGTTAATATTTTGTTAAAATTCGCGTT |

| | |
|---|---|
| | AAATTTTTGTTAAATCAGCTCATTTTTTAACGAATAGCCCGAAATCGGCAAAATCCCTTA<br>TAAATCAAAAGAATAGACCGAGATAGGGTTGAGTGTTGTTCCAGTTTCCAACAAGAGTCC<br>ACTATTAAAGAACGTGGACTCCAACGTCAAAGGGCGAAAAAGGGTCTATCAGGGCGATGG<br>CCCACTACGTGAACCATCACCCTAATCAAGTTTTTTGGGGTCGAGGTGCCGTAAAGCAGT<br>AAATCGGAAGGGTAAACGGATGCCCCCATTTAGAGCTTGACGGGGAAAGCCGGCGAACGT<br>GGCGAGAAAGGAAGGGAAGAAAGCGAAAGGAGCGGGGGCTAGGGCGGTGGGAAGTGTAGG<br>GGTCACGCTGGGCGTAACCACCACACCCGCCGCGCTTAATGGGGCGCTACAGGGCGCGTG<br>GGGATGATCCACTAGTACGGATTAGAAGCCGCCGAGCGGGTGACAGCCCTCCGAAGGAAG<br>ACTCTCCTCCGTGCGTCCTCGTCCTCACCGGTCGCGTTCCTGAAACGCAGATGTGCCTCG<br>CGCCGCACTGCTCCGAACAATAAAGATTCTACAATACTAGCTTTTATGGTTATGAAGAGG<br>AAAAATTGGCAGTAACCTGGCCCCACAAACCTTCAAATGAACGAATCAAATTAACAACCA<br>TAGGATGATAATGCGATTAGTTTTTTAGCCTTATTTCTGGGGTAATTAATCAGCGAAGCG<br>ATGATTTTTGATCTATTAACAGATATATAAATGCAAAAACTGCATAACCACTTTAACTAA<br>TACTTTCAACATTTTCGGTTTGTATTACTTCTTATTCAAATGTAATAAAAGTATCAACAA<br>AAAATTGTTAATATACCTCTATACTTTAACGTCAAGGAGAAAAAACCCCGGATCGGACTA<br>CTAGCAGCTGTAATACGACTCACTATAGGGAATATTAAGCTTGGTACCctcgagATGGAT<br>GTATTCATGAAAGGACTTTCAAAGGCCAAGGAGGGAGTTGTGGCTGCTGCTGAGAAAACC<br>AAACAGGGTGTGGCAGAAGCAGCAGGAAAGACAAAAGAGGGTGTTCTCTATGTAGGCTCC<br>AAAACCAAGGAGGGAGTGGTGCATGGTGTGGCAACAGTGGCTGAGAAGACCAAAGAGCAA<br>GTGACAAATGTTGGAGGAGCAGTGGTGACGGGTGTGACAGCAGTAGCCCAGAAGACAGTG<br>GAGGGAGCAGGGAGCATTGCAGCAGCCACTGGCTTTGTCAAAAAGGACCAGTTGGGCAAG<br>AATGAAGAAGGAGCCCCACAGGAAGGAATTCTGGAAGATATGCCTGTGGATCCTGACAAT<br>GAGGCTTATGAAATGCCTTCTGAGGAAGGGTATCAAGACTACGAACCTGAAGCCGGCGCC<br>CAGATCCACCGGTCGCCAACCGCAGTGAGCAAGGGCGAGGAGCTGTTCACCGGGGTGGTG<br>CCCATCCTGGTCGAGCTGGACGGCGACGTAAACGGCCACAAGTTCAGCGTCCGCGGCGAG<br>GGCGAGGGCGATGCCACCAACGGCAAGCTGACCCTGAAGTTCATCTGCACCACCGGCAAG<br>CTGCCCGTGCCCTGGCCCACCCTCGTGACCACCTTCGGCTACGGCGTGGCCTGCTTCAGC<br>CGCTACCCCGACCACATGAAGCAGCACGACTTCTTCAAGTCCGCCATGCCCGAAGGCTAC<br>GTCCAGGAGCGCACCATCTCTTTCAAGGACGACGGTACCTACAAGACCCGCGCCGAGGTG<br>AAGTTCGAGGGCGACACCCTGGTGAACCGCATCGAGCTGAAGGGCATCGACTTCAAGGAG<br>GACGGCAACATCCTGGGGCACAAGCTGGAGTACAACTTCAACAGCCACAACGTCTATATC<br>ACGGCCGACAAGCAGAAGAACGGCATCAAGGCTAACTTCAAGATCCGCCACAACGTTGAG<br>GACGGCAGCGTGCAGCTCGCCGACCACTACCAGCAGAACACCCCCATCGGCGACGGCCCC<br>GTGCTGCTGCCCGACAACCACTACCTGAGCCATCAGTCCGCCCTGAGCAAAGACCCCAAC<br>GAGAAGCGCGATCACATGGTCCTGCTGGAGTTCGTGACCGCCGCCGGGATTACACATGGC<br>ATGGACGAGCTGTACAAGTAANNNNNNNNNNNNNNNNNNNNNNNNNNgagctcTCTAGAG<br>GGCCGCATCATGTAATTAGTTATGTCACGCTTACATTCACGCCCTCCCCCCACATCCGCT<br>CTAACCGAAAAGGAAGGAGTTAGACAACCTGAAGTCTAGGTCCCTATTTATTTTTTTATA<br>GTTATGTTAGTATTAAGAACGTTATTTATATTTCAAATTTTTCTTTTTTTTCTGTACAGA<br>CGCGTGTACGCATGTAACATTATACTGAAAACCTTGCTTGAGAAGGTTTTGGGACGCTCG<br>AAGGCTTTAATTTGCGGCC |

**Table S6.** Fitness scores of familial Parkinson's disease variants at different protein expression levels

| | Fitness Scores at | | | |
|---|---|---|---|---|
| Familiar PD variant | 0.001% | 0.01% | 0.1% | 1% |
| A30P | 0.0841 | 0.152 | 0.5345 | 0.5500 |
| E46K | -0.0304 | -0.0168 | 0.0175 | 0.0488 |
| H50Q | -0.0446 | -0.0192 | -0.0296 | -0.0262 |
| G51D | 0.0254 | 0.1483 | 0.2512 | 0.2988 |
| A53T | -0.0219 | 0.0422 | 0.063 | 0.0126 |
